# Genomic context sensitivity of insulator function

**DOI:** 10.1101/2021.05.16.444030

**Authors:** André M. Ribeiro-dos-Santos, Megan S. Hogan, Raven D. Luther, Matthew T. Maurano

## Abstract

Compartmentalization of interactions between genomic regulatory elements and potential target genes is influenced by the binding of insulator proteins such as CTCF, which act as potent enhancer blockers when interposed between an enhancer and a promoter in a reporter assay. But only a minority of CTCF sites genome-wide function as boundary elements, depending on cellular and genomic context. To dissect the influence of genomic context on enhancer blocker activity, we integrated reporter constructs with promoter-only, promoter and enhancer, and enhancer blocker configurations at hundreds of thousands of genomic sites using the Sleeping Beauty transposase. Deconvolution of reporter activity by genomic position revealed strikingly different patterns of reporter function, including a compartment of enhancer blocker reporter integrations with robust expression. The high density of integration sites permits quantitative delineation of characteristic genomic context sensitivity profiles, and their decomposition into sensitivity to both local and distant DNaseI hypersensitive sites. Furthermore, a single-cell expression approach permits direct linkage of reporters integrated into the same clonal lineage with differential endogenous gene expression, revealing that CTCF insulator activity does not completely abrogate reporter effects on endogenous gene expression. Collectively, our results lend new insight to genomic regulatory compartmentalization and its influence on the determinants of promoter-enhancer specificity.

## Introduction

Insulators are a class of genomic regulatory elements that block interaction of enhancers with their cognate promoters (Phillips and Corces 2009). A strict boundary model offers an attractive paradigm for understanding regulatory specificity in mammalian genomes through the delineation of regulatory domains. Enhancer blocker activity is canonically defined by a reporter assay which interposes a candidate insulator element between a weak promoter and an enhancer (Chung et al. 1993), while barrier insulators protect transgenes from silencing due to spreading of heterochromatin (West et al. 2002). Insulators have also been employed to counter genotoxicity from transgene enhancer activation of endogenous oncogenes (Li et al. 2009; Liu et al. 2015). Known insulators such as the chicken ß-globin hypersensitive site 4 element or the Igf2/H19 imprinting control region (Bell and Felsenfeld 2000) are composite elements with enhancer blocker, barrier, and other activities (Dickson et al. 2010), and often have secondary functions, such as silencers (Qi et al. 2015).

The architectural protein CTCF is the only known vertebrate insulator protein and its binding can confer a potent enhancer blocking effect (Phillips and Corces 2009). Additionally, binding sites for CTCF co-localize with genomic features such as topologically associated domain boundaries (Dixon et al. 2012), but direct functional analysis of these sites is impeded by the difficulty of genome engineering at the relevant scales. While binding affinity and recognition sequence orientation appear to confer some specificity for CTCF sites involved in domain organization (Guo et al. 2015; de Wit et al. 2015), these factors alone remain inadequate to distinguish true barrier elements from the ∼100,000 cell-type specific CTCF sites genome-wide (Maurano et al. 2015). Stably integrated reporter assays have shed light on the mechanics of insulator function, but such methods do not assess interaction with the surrounding endogenous genomic elements (Walters et al. 1999). In contrast, integrated barcoded reporter assays (Akhtar et al. 2013; Maricque et al. 2018; Moudgil et al. 2020) offer the potential to directly assess the interaction between novel CTCF sites and the endogenous genomic landscape.

Here we describe a high-throughput, randomly integrated barcoded reporter platform to analyze insulator activity in varied genomic contexts. We developed an enhancer blocker construct interposing a potent CTCF insulator element (Liu et al. 2015) between a ß-globin HS2 enhancer (HS2) and Ɣ-globin promoter. Barcoded reporters with or without insulator elements were randomly integrated into the genome of cultured K562 cells using the Sleeping Beauty transposase system, and subsequently mapped to enable barcode-specific readout of genomic context effects. We find that reporters are distinguished by characteristic response signatures to genomic context, modulated by the presence of enhancer and insulator elements. Finally, we employ single cell RNA-seq to link cells deriving from the same initial clone and link specific integrations to perturbation of endogenous gene expression.

## Results

### Flow cytometry characterization of enhancer blocker reporter

We developed and characterized a series of reporter constructs based on well-characterized genomic regulatory elements including the murine Ɣ-globin promoter and murine ß-globin locus control region (LCR) hypersensitive site 2 (HS2) enhancer. A potent insulator element (A1 or C1) previously identified through an analysis of highly occupied CTCF sites in the human genome (Liu et al. 2015) (**Supplemental Fig. S1**) was interposed between the promoter and enhancer in an enhancer blocker position (**Fig. 1**). Reporter expression drove a PuroGFP fusion protein to enable selection and/or measurement of transcriptional activity on a cellular level. The reporter was flanked by Sleeping Beauty (Mátés et al. 2009) inverted terminal repeats (ITRs) to enable transposition into the genome. Reporter plasmids were stably integrated at random genomic sites through transient co-transfection with a plasmid expressing SB100X, a highly active variant of the Sleeping Beauty transposase (Mátés et al. 2009).

**Fig. 1.**
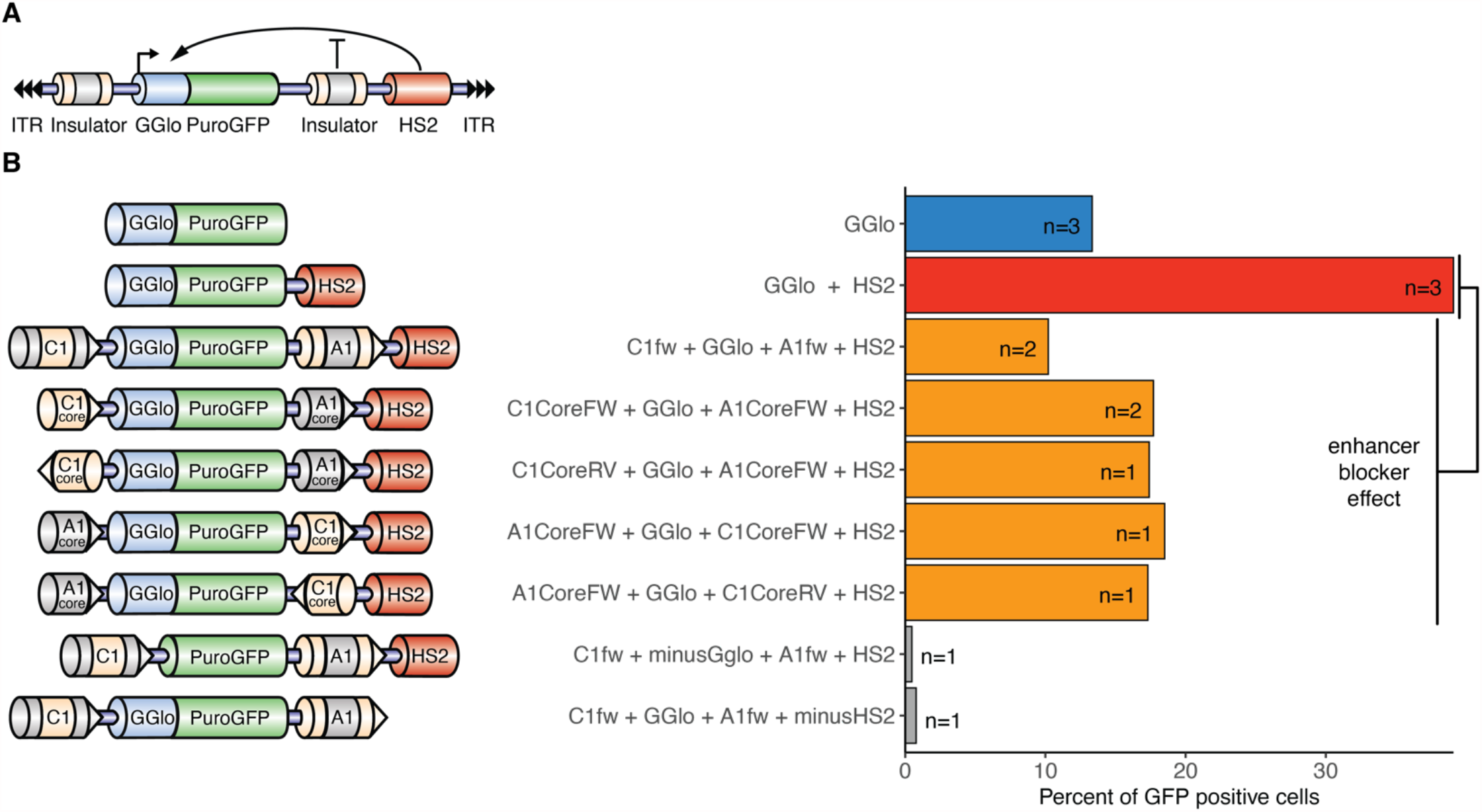
Cellular activity of enhancer blocker activity. (A) Reporter scheme consisting of Ɣ-globin promoter driving PuroGFP expression. An intervening CTCF site between the promoter and an HS2 enhancer acts as an enhancer blocker. ITR, Sleeping Beauty inverted terminal repeats; GGlo, Ɣ-globin promoter; HS2, ß-globin hypersensitive site 2 enhancer. (B) Reporter plasmids were transfected genomically integrated by co-transfection with a plasmid expressing the Sleeping Beauty SB100X transposase. Activity was measured by flow cytometry. A1 and C1 represent previously characterized CTCF-binding insulator elements (Liu et al. 2015).

We first characterized the activity of several different classes of reporters based on this scaffold, including GGlo (promoter-only), GGlo+HS2 (promoter and enhancer), and Ins+GGlo+Ins+HS2 (enhancer blocker) reporters. K562 erythroleukemia cells were transfected with both reporter plasmid and SB100X transposase plasmid (**Supplemental Table S1**). Reporter activity was characterized using flow cytometry to measure the proportion of GFP+ cells (**Fig. 1, Supplemental Fig. S2**). Ins+GGlo+Ins+HS2 reporters showed low GFP+ cells, comparable to promoter-only GGlo constructs, confirming that the CTCF site acts as an enhancer blocker. An insulator element truncated to the core 54 bp of the CTCF recognition sequence (**Supplemental Fig. S1)** similarly showed lowered GFP+ cells (**Fig. 1**), confirming that the enhancer blocker effect is tightly coupled to the CTCF recognition sequence. Reporters without a promoter or enhancer showed essentially no GFP+ cells, confirming that neither the insulator elements nor Sleeping Beauty ITRs have intrinsic transcriptional or enhancer activity on their own. Finally, CTCF effect on reporter activity was orientation-independent. These results confirm that our reporter assay detects canonical enhancer blocker reporter activity.

### Reporter activity in genomic context

Flow cytometry assesses single-cell GFP activity representing the sum of all reporters integrated in that cell, but individual insertion sites may exhibit a wide range of activity. We developed a strategy to deconvolute the activity of individual reporters using unique 16 nt reporter barcode (BC) sequences. Sequencing libraries were constructed using a two-stage nested PCR to add Illumina adapters. We generated three types of libraries based on this reporter BC strategy (**Fig. 2A, Supplemental Fig. S3, Supplemental Fig. S4**): reporter genomic integration sites were mapped using inverse PCR (iPCR), their representation was determined using DNA libraries, and their expression was measured with RNA libraries. RNA and DNA libraries incorporated a 8-12 nt unique molecular identifier (UMI) (Jee et al. 2016) to permit targeted single-molecule counting.

**Fig. 2.**
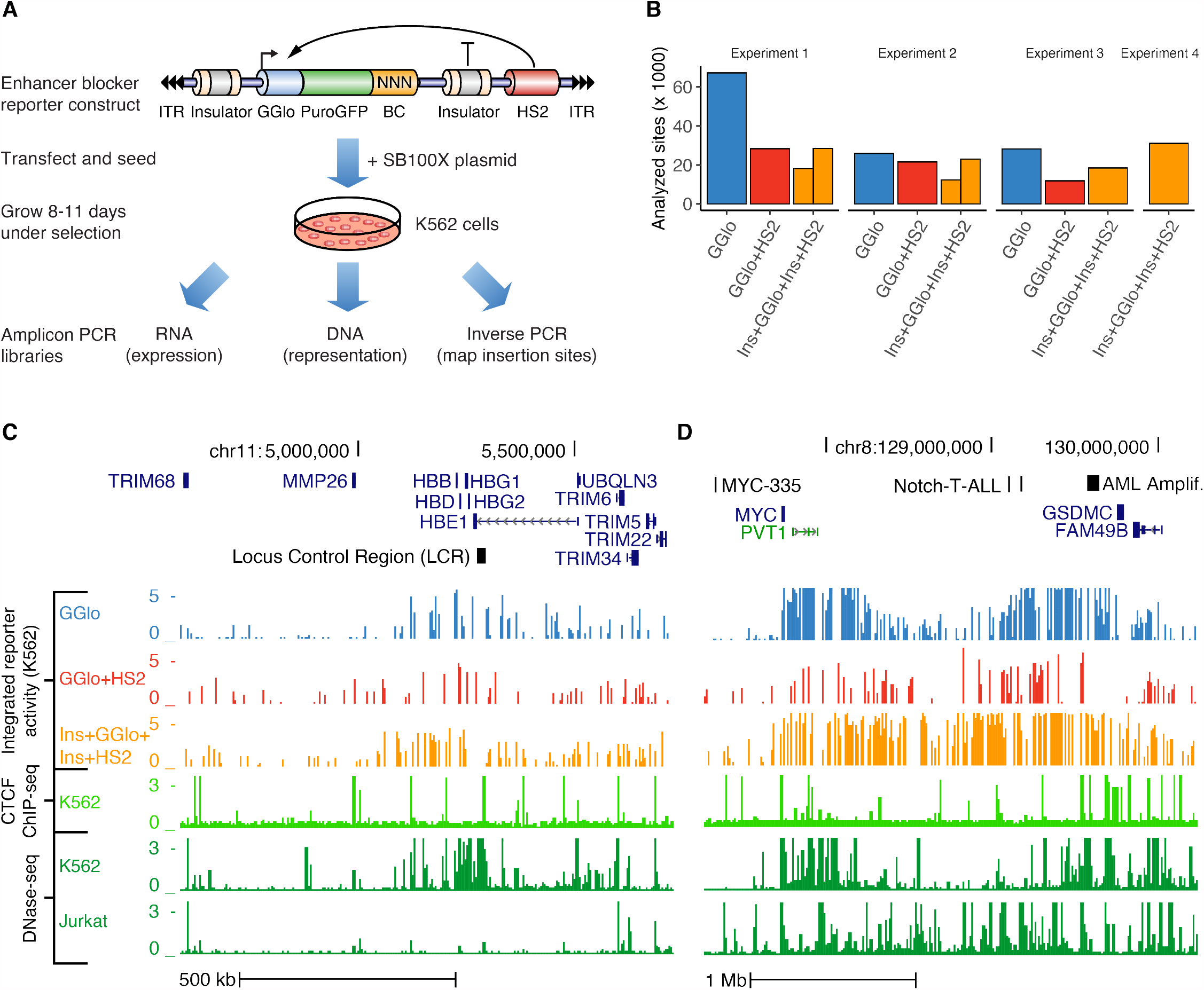
Site-specific analysis of enhancer blocker activity. (A) To measure site-specific reporter activity, barcoded reporter plasmids are randomly integrated through co-transfection with a plasmid expressing SB100X transposase and read out in multiplex. PCR-based Illumina library construction enables highly quantitative measurement of hundreds of thousands of barcodes simultaneously. Insertion sites are mapped using inverse PCR; DNA libraries are used to normalize for barcode representation, and RNA libraries to quantify barcode expression. GGlo, Ɣ-globin promoter; BC, unique barcode; HS2, B-globin hypersensitive site 2 enhancer; ITR, Sleeping Beauty inverted terminal repeats. (B) Counts of sites analyzed for 4 replicate experiments of reporters containing promoter only (GGlo), promoter and HS2 enhancer (GGlo+HS2), or with CTCF site interposed between GGlo and HS2 (Ins+GGlo+Ins+HS2). (C-D) Analysis of enhancer-blocker functionality at the *HBB* (C) and *MYC* (D) loci. Top tracks show reporter activity. Shown are data merged from all replicate experiments. Bottom tracks show CTCF ChIP-seq data for K562 erythroleukemia cells, and DNase-seq data for K562 and Jurkat T-cell leukemia cells. Regions highlighted in (D) include: the MYC-335 enhancer region coinciding with a genetic association for colorectal cancer (Sur et al. 2012), the Notch-T-ALL (Acute Lymphocytic Leukemia) enhancer cluster (Herranz et al. 2014), and the AML (acute myeloid leukemia) amplified region (Radtke et al. 2009).

We performed a series of 4 replicate experiments using these GGlo, GGlo+HS2, and Ins+GGlo+Ins+HS2 constructs (**Supplemental Table S1**). After growth under puromycin selection for 8-11 days, multiple DNA, RNA, and iPCR libraries (n = 2-4) were generated for each experiment (**Supplemental Table S2**) and sequenced to saturation. Reporter activity was established as the log-ratio of RNA and DNA counts. Individual transfections averaged 31,529 measured reporter insertions analyzed after quality control (**Fig. 2B, Supplemental Table S3**). This activity recapitulated cellular activity when averaged across all insertion sites (**Fig. 1, Supplemental Fig. S5A**). These data yielded high-resolution maps of reporter activity, with a reporter insertion every 23-48 kb on average, and an average distance from an endogenous DNaseI hypersensitive site (DHS) to a reporter insertion of 9-19 kb.

Replicate libraries from the 4 experiments were merged together to maximize resolution for visualization and analysis, yielding 314,796 insertion sites in total (**Table 1**; **Supplemental Table S3**). Examination of the ß-globin (**Fig. 2C**) and *MYC* (**Fig. 2D**) loci demonstrated notable differences in patterns of reporter activity. GGlo exhibited variable activity that was highly responsive to local genomic context: at the ß-globin locus, its activity was concentrated tightly at genes; at *MYC*, activity localized to two separate domains around the *MYC* gene itself and distal ALL enhancers. GGlo+HS2 showed more variable insertion location and site-specific activity. We observed peak activity at a subset of regions, but insertions were depleted over several regions, including the DHS cluster immediately downstream of *MYC*, suggesting that the GGlo+HS2 construct does not support expression at these regions, or that those insertions have a negative effect on growth. Ins+GGlo+Ins+HS2 exhibited more uniform activity regardless of surrounding genomic context and showed reduced position preference throughout the window, consistent with the notion that CTCF sites moderate the impact of surrounding genomic context on regulatory activity.

**Table 1.** Summary of insertions analyzed per reporter class and experiment. Summary of transfections across four independent experiments. Counts are of insertions passing all QC filters.

| Reporter Class | Experiment 1 | Experiment 2 | Experiment 3 | Experiment 4 | Merged |
| --- | --- | --- | --- | --- | --- |
| GGlo | 66,742 | 25,930 | 28,107 | - | 120,604 |
| GGlo+HS2 | 28,305 | 21,572 | 11,974 | - | 61,796 |
| Ins+GGlo+Ins+HS2 | 46,566 | 35,348 | 18,469 | 30,935 | 131,073 |

To systematically assess the length scale of sensitivity of different reporters to genomic context, we computed the correlation in activity for insertions at adjacent sites (**Fig. 3A, Supplemental Fig. S5B**). GGlo and Ins+GGlo+Ins+HS2 showed high correlation at short-range (<5 kb), suggesting that local genomic context played a significant role in reporter activity. However, at longer range, GGlo correlation dropped to nearly zero while Ins+GGlo+Ins+HS2 remained high beyond 50 kb. GGlo+HS2 showed the lowest correlation across all size ranges, suggesting the least influence of genomic context.

**Fig. 3.**
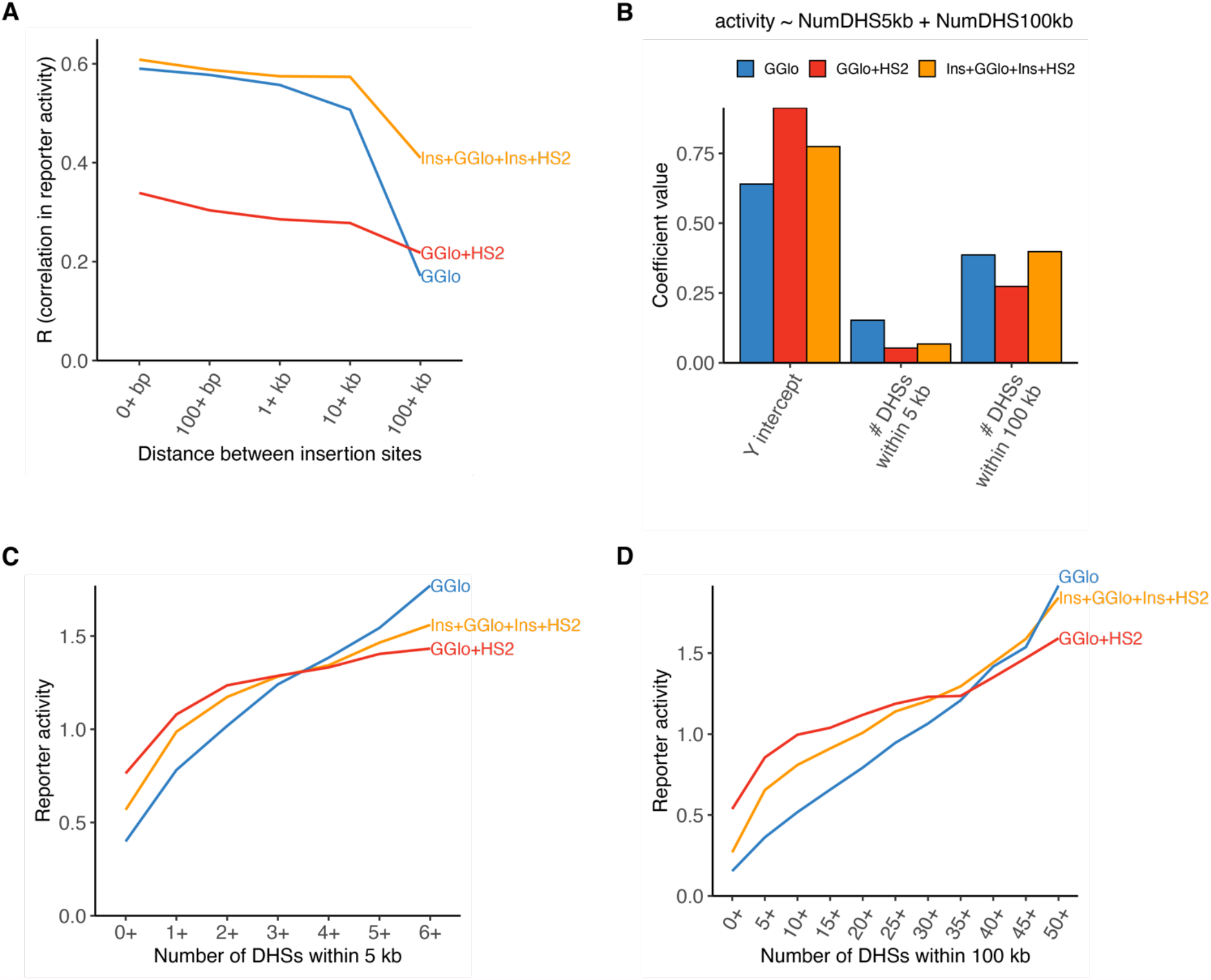
Quantitative assessment of genomic context effects on reporter activity. (A) Correlation in activity for nearby insertions by reporter class. Data is merged across all experiments; individual experiments are shown in **Supplemental Fig. S5B**. (B) Linear regression coefficients for regression of reporter activity on density of local and long-range genomic context. (C-D) Reporter activity by short-range and long-range genomic context represented by the number of DHSs within 5 kb (C) or 100 kb (D).

To provide an easily computed metric reflecting the contribution of genomic context at different distance scales, we computed the number of DHSs within 5 kb and 100 kb of the reporter insertion site. We then used a regression linear model to systematically quantify the effect of these indicators of genomic context on activity of reporters of different classes (**Fig. 3B**). Consistent with the correlation pattern observed at nearby insertions (**Fig. 3A**), GGlo and Ins+GGlo+Ins+HS2 showed the highest contribution of genomic context factors to reporter activity (R^2^ = 0.28 and 0.27, respectively, while GGlo+HS2 showed the lowest (R^2^ = 0.16). All three reporters showed distinct contributions of short and long-range genomic context (**Fig. 3C-D**). The relevance of DHS density as a feature extending as far as 100 kb suggests an infinitesimal model where genomic context reflects the influence of a large number of regulatory elements, with most contributing a small but significant effect.

### Clonal analysis using integrated barcodes

Single-cell RNA-seq (scRNA-seq) approaches can provide the compartmentalization needed to associate reporter BCs integrated in the same cell, which can provide a unique combinatorial genetic identifier for cells derived from a given clone during transfection (Lu et al. 2011; Biddy et al. 2018; Weinreb et al. 2020). We adapted our integrated reporter assay to the droplet-based 10x Genomics scRNA-seq platform and performed a pilot experiment (Experiment 4) using Ins+GGlo+Ins+HS2 (**Fig. 4A**; **Supplemental Table S4**). We used an amplicon-targeted library construction approach to enrich scRNA-seq cDNA for reporter transcripts. These data showed that reporter activity was highly reproducible across multiple cells. For comparison, control comparisons of different reporters in the same cells or with permuted data showed little correlation (**Fig. 4B**).

**Fig. 4.**
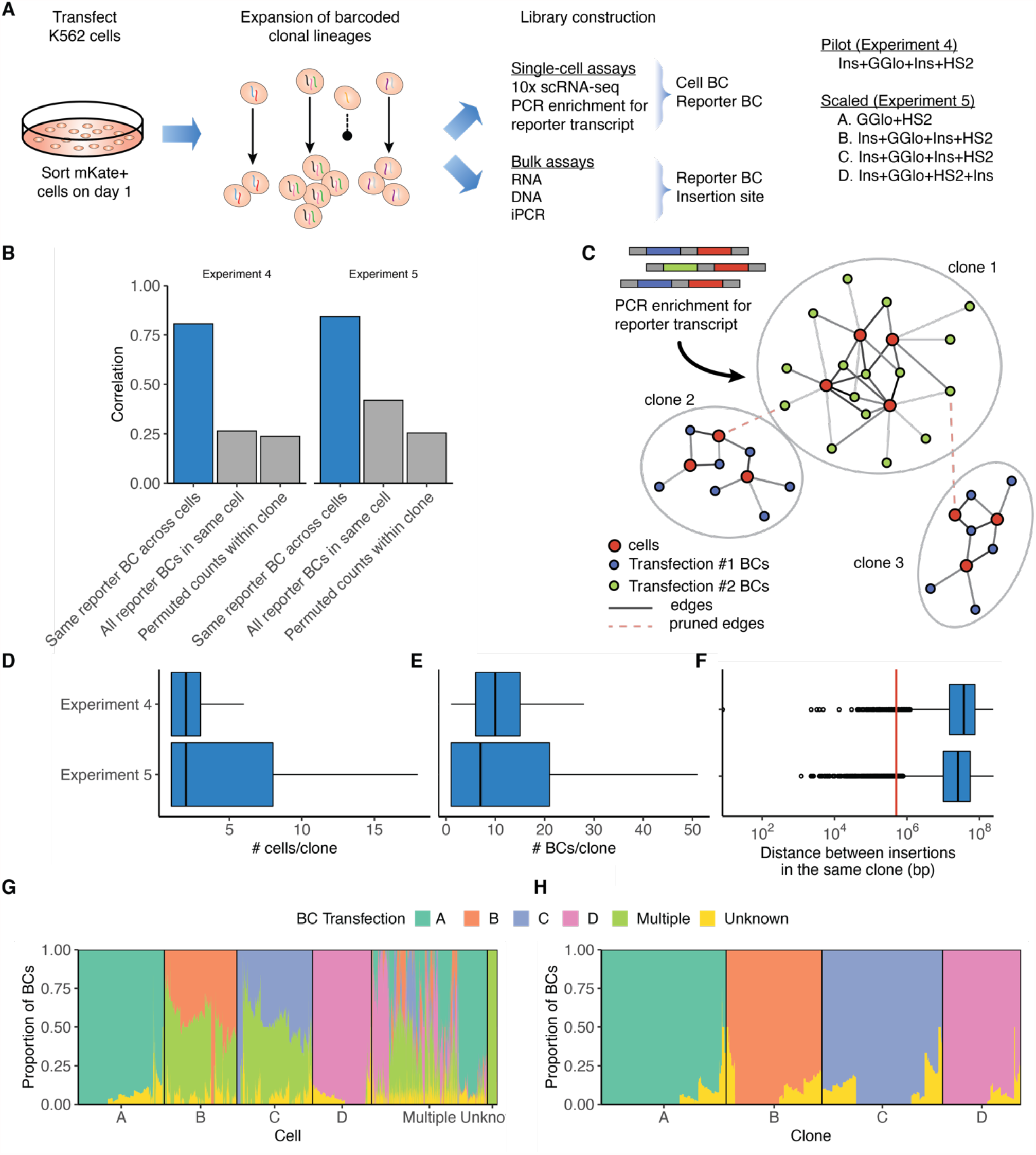
scRNA-seq inference of clonal relationship of reporter insertions. (A) scRNA-seq experiment to infer clonal relationships between single cells using presence of reporter BCs. (B) Reporter BC counts show high correlation across multiple cells. All reporter BCs in the same cells or reporter BC counts shuffled within clones show little correlation. (C) Graph-based inference of clones from scRNA-seq data using reporter BC to link cells deriving from the same clone. (D-F) Summary of clones including Number of cells per clone (D), number of reporter BCs per clone (E), and distance between reporters integrated in cells derived from the same clone (F). Boxes represent quartiles, vertical bar indicates median, and whiskers represent ±1.5 interquartile range. Vertical red line in (F) at 500 kb indicates insertions considered far enough to be assumed independent. (G-H) Deconvolution of Experiment 5. X-axis represents starting cells (G) or final inferred clones (H), grouped by inferred transfection and ordered according to hierarchical clustering. Multiple, reporter BCs detected in multiple transfections; Unknown, reporter BCs detected only in scRNA-seq data.

We developed a graph-based clonal inference approach to identify cells deriving from the same clone. The Experiment 4 data was used to initialize a graph by defining cellBCs and reporter BCs as nodes which were connected based on the reporter-enriched scRNA-seq libraries (**Fig. 4C**). The graph was then pruned to reduce the impact of typical sources of experimental noise in large screens, such as chimeric PCR products and cell doublets (Methods). From an original set of 3,886 cells, we identified 1,832 clones (**Supplemental Data S1**). Clones contained a median of 2 cells (**Fig. 4D**) and a median of 10 reporter BCs (**Fig. 4E**). Assessing reporter insertion sites in the same cell showed that the distance between insertions was sufficiently large to enable independent readout of hundreds or thousands of reporters in a given cell (**Fig. 4F**).

Based on these results, we performed a scaled-up experiment (Experiment 5) using 3 different classes of reporter constructs, including GGlo+HS2, Ins+GGlo+Ins+HS2 (transfected in replicate from the same plasmid library), and Ins+GGlo+HS2+Ins (a reporter fully flanked by insulator elements). Given that each transfection is distinguished by a distinct set of reporter BCs, we pooled the cells from four independent transfections. We generated 4 replicate 10x scRNA-seq libraries, all of which were superloaded to maximize power (**Supplemental Table S4)**. Reporter BCs specific to each individual transfection were identified using the DNA, RNA, and iPCR libraries from bulk cells. The majority of cells were assignable to a single transfection, with the exception of reporter BCs shared from the replicate transfections B and C (**Fig. 4G**). We extended our clonal inference approach to used Reporter BC labels to prune conflicting cells and clones (**Fig. 4H**). The final results include a total of 4,576 clones (**Supplemental Data S2)**.

### scRNA-seq differential expression analysis

The compartmentalization provided by single-cell readout enables direct linkage of specific insertions to their effect on adjacent genes (Gasperini et al. 2019). To investigate the effect of ectopic regulatory DNA on the local genomic landscape, we developed an analysis approach called Differential Expression of Clonal Alterations Local effects (DECAL) which identifies nearby genes with significant differential expression while accounting for the sparsity of single-cell data (**Fig. 5A, Methods**). Use of clonal inference permitted imputation across cells of reporter presence and flanking gene expression levels, and substantially increased the number of cells linked to a particular reporter beyond those with direct measurement of the reporter BC (**Supplemental Fig. S6A**). Expression levels were highly consistent between cells with direct measurement of the reporter BC and other cells in the same clone (**Supplemental Fig. S6B**).

**Fig. 5.**
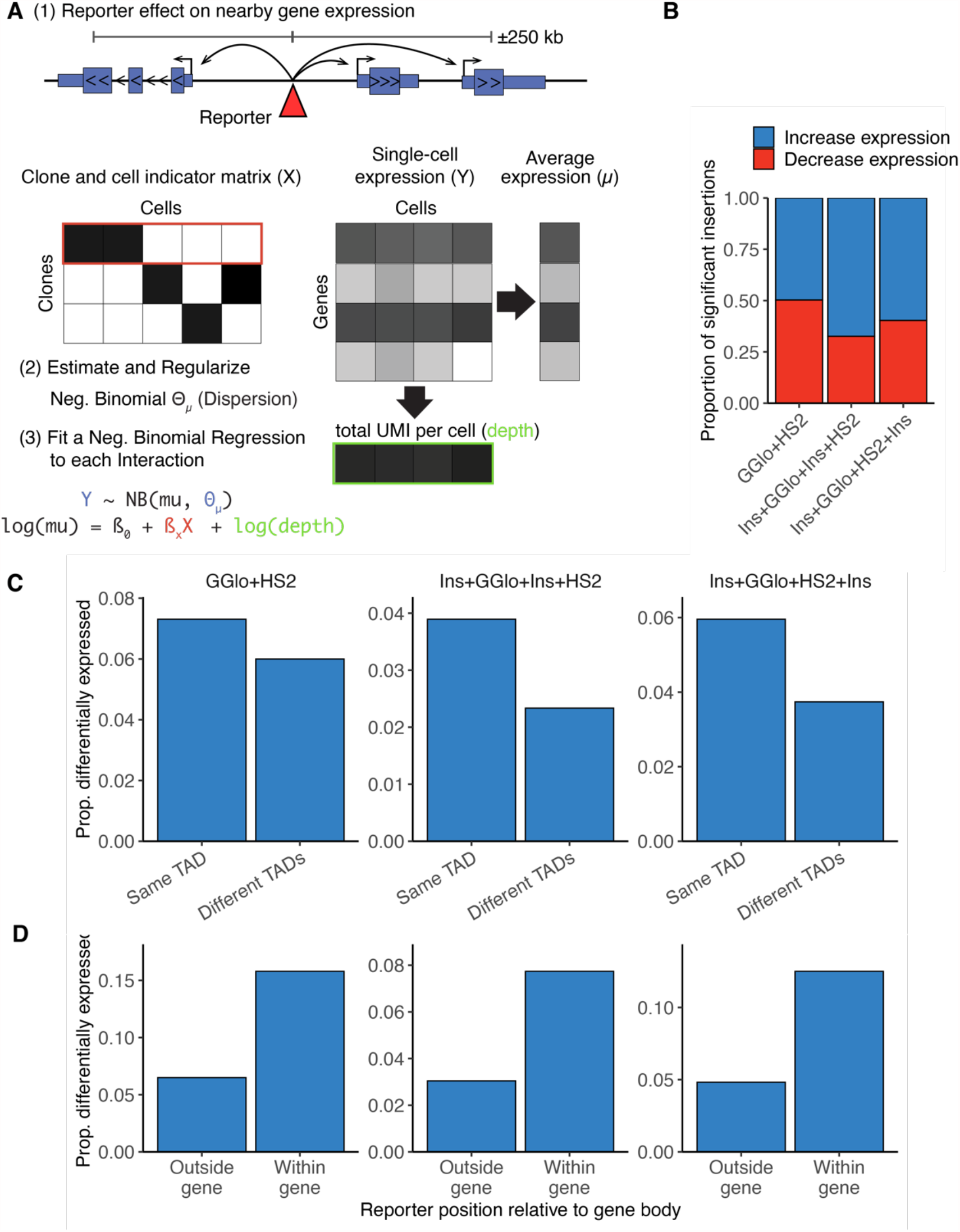
Analysis of reporter effect on nearby endogenous gene expression. (A) Schematic of analysis framework. (B) Significant tests broken down into those that increase vs. decrease expression. (C-D) Rate of significant effect on gene expression by (C) whether reporter and gene are in the same TAD; and (D) whether reporter is inside or outside gene body.

We then applied DECAL to analyze the full-scale data in Experiment 5. We first conducted a power simulation to estimate the ability of our statistical framework to detect expression perturbations (**Supplemental Fig. S7**). Using these results, we restricted tests to those well powered to detect expression changes (**Methods**). We then tested for significant effects on gene expression for each of the three distinct reporter classes, pooling results from the replicate Ins+GGlo+Ins+HS2. There was sufficient power to test an average of 3,122 insertions per reporter class (**Table 2, Supplemental Data S3**). While most reporters did not have an effect on nearby endogenous gene expression, a small proportion (3.2%-6.8%) of reporters of all three classes caused detectable effects on expression of a nearby gene (**Table 2**). The direction of effect for significant insertion-gene tests was evenly split between increased and decreased expression of the target gene (**Fig. 5B**). Insertions were more likely to affect gene expression of TSSs in the same TAD (**Fig. 5C**) and when the reporter was inserted in the gene body itself (**Fig. 5D**). Collectively, these results underscore the role for genomic context in dictating the effect of ectopically delivered regulatory elements on endogenous gene expression.

**Table 2.** Effect of reporter insertion on endogenous gene expression. Genes within 250 kb of a reporter insertion were tested for differential expression. Summary of the number of tests, unique reporters, unique genes, and significant tests (q-value < 0.05) by reporter class.

| Reporter Class | # tests | # reporters tested | # genes tested | # significant | Avg. distance to TSS of significant tests |
| --- | --- | --- | --- | --- | --- |
| GGlo+HS2 | 2,808 | 2,110 | 868 | 193 | 122 kb |
| Ins+GGlo+Ins+HS2 | 4,174 | 3,079 | 989 | 135 | 124 kb |
| Ins+GGlo+HS2+Ins | 2,303 | 1,711 | 876 | 119 | 122 kb |

## Discussion

While present day classes of genomic regulatory elements were originally defined using reporter assays, short constructs alone cannot model the effect of hundreds or thousands of regulatory sites which collectively influence genomic context. Here we show that while insulator elements demonstrate a strong effect in aggregate, individual reporters demonstrate a wide range of expression levels depending on genomic context. Indeed, the high expression at some genomic integration sites implies a total abrogation of insulator function. Similarly, integrated reporters perturbed endogenous gene expression regardless of the presence of insulating CTCF sites. A strict barrier definition of insulator function is difficult to reconcile with these results, or with the abundance of CTCF sites in the genome.

Our results are instead consistent with a model in which insulators moderate but do not eliminate sensitivity to genomic context. Single-cell analyses have shown that enhancers increase the frequency of transcription in a given cell rather than augmenting the transcriptional rate per cell (Weintraub 1988; Walters et al. 1996). Our data show that independent but nearby (<1 kb) insertions of GGlo and Ins+GGlo+Ins+HS2 reporters exhibit high correlation in expression; GGlo+HS2 reporter expression showed a much lower correlation at nearby sites, suggesting a higher influence of epigenetic or stochastic factors (**Supplemental Fig. S5B**). These results suggest that insulator elements might counteract increased stochasticity from an enhancer element while still leaving expression subject to position effects. It is possible that more complex, composite regulatory elements might further fine tune variability of expression and sensitivity to context effects. For example, multiple tandem CTCF sites have been shown to increase the strength of insulation (Huang et al. 2021).

Genomic context is a key predictive feature of models for recognizing functional regulatory variation (Halow et al. 2021), suggesting a need for further large-scale characterization of genomic regulatory element classes and composite regulatory elements. Our work suggests that genomic regulatory element function should be evaluated on multiple axes, including expression (i) level, (ii) consistency (stochastic vs. deterministic), and (iii) sensitivity to the local and/or long-range regulatory landscape. We expect that our approach will enable further dissection of the interplay between regulatory sequence, genomic context, and single-cell behavior and facilitate incorporation of genomic context sensitivity in models of functional regulatory variation

## Methods

### Plasmid cloning and barcoding

pCMV(CAT)T7-SB100 (SB100X) and pT2/LTR7-GFP were gifts from Zsuzsanna Izsvak (Addgene plasmids #34879 and #62541, respectively) (Mátés et al. 2009).

The Ɣ-globin promoter, mouse ß-globin hypersensitive site 2 (HS2) enhancer, A1 insulator, A2 insulator, C1 insulator, A1 Core, and C1 Core DNA fragments (**Supplemental Table S5**) were synthesized by Genscript USA (Piscataway, NJ). All plasmids used in this study are listed in **Supplemental Table S6**.

Sleeping Beauty reporter constructs used in this study were barcoded using a Gibson Assembly approach prior to introduction into K562 cells (**Supplemental Fig. S3**). The plasmid backbone to be barcoded was PCR amplified using pTR-GibsonBC-FW and pTR-GibsonBC-RV primers, and the correct length fragment was purified from a 1% agarose gel. Next, Gibson Assembly was performed using the amplified plasmid backbone and a synthesized DNA fragment “GibsonBC4” according to the manufacturer’s protocol (NEB cat# E2611L). Barcoded plasmid library DNA was purified using the Zymo Clean and Concentrate-5 (Zymo Research cat# D4014) protocol prior to transformation. Purified barcoded plasmid DNA was transformed into electrocompetent MegaX DH10B-T1 bacteria (Fisher cat# C640003) using an Eppendorf 2510 electroporator set to 1800 V. After recovering for 1 h at 37 °C, transformation reactions were transferred to 50 mL LB Media with 100 µg/mL Ampicillin and incubated at 37 °C for 16 h shaking at 220 RPM. Barcoded plasmid library DNA was purified using the ZymoPure II Plasmid Maxiprep kit protocol and quantified on a Nanodrop.

### Cell culture and transfection

K562 cells were obtained from ATCC (ATCC CCL-243) and cultured in RPMI 1640 medium with glutamine (Fisher cat# MT10040CV) supplemented with 10% FBS (Gemini Bio-Products cat# 100-106), 1 mM sodium pyruvate, and 10 U/mL penicillin-streptomycin. Cultures were maintained at 37 °C and 5% CO_2_, and were subcultured once cultures reached a density of 5×10^5^ cells/mL.

×10^6^ K562 cells were transfected using the ThermoFisher Neon Transfection System 100 μL Kit according to the manufacturer’s instructions with varying amounts of transposon and transposase (**Supplemental Table S1**). Cells were transfected with 4 μL TE to use as a negative control for puromycin selection. Transfected cells were selected with puromycin (2.5 µg/mL). K562 media with puromycin was replaced every 2 days. Cell counts were performed either using PrestoBlue (Life Technologies cat# A13261) and fluorescence detection with the Synergy H1 Multi-Mode Microplate Reader, or were stained with trypan blue and counted on a hemocytometer.

### Flow cytometry of GFP Expression Assays

On day 8 after transfection, GFP expression was measured using the Sony SH800S Cell Sorter. For each experiment, a 100 µM chip and the Optical Filter Pattern 2 were used, the 405 nm, 488 nm and 561 nm lasers were enabled, automatic color compensation was turned off, and sensor gain settings were set to the following values: forward scatter (FSC) = 3, back scatter (BSC) = 30.5%, and FL2 (GFP) = 36.5%.

Using FlowJo software (v10.7.2), single live cells were gated using side scatter (SSC) and FSC values from a TE (mock) transfected cell sample. GFP expression data were plotted on a histogram of unit area vs. GFP fluorescence (525±50 nm). The GFP-negative cell population was defined using the GFP expression of TE (mock) transfected single-cells. The GFP-positive cell population was defined by cells that had a greater GFP fluorescence than TE (mock) transfected cells.

### Genomic DNA Purification

11-14 days post-transfection, cell pellets containing 3×10^6^ to 4×10^6^ cells each were snap frozen in LN_2_ and stored at −80°C until DNA extraction. Cell pellets were allowed to warm to room temperature, and then were resuspended in 385 µL DNA Quick Extract (Lucigen cat# QE09050) and transferred to a 1.5 mL tube. Cells were incubated at 65 °C for 15 min, followed by 98 °C for 5 min. After cooling briefly, 10 µL Proteinase K (Sigma-Aldrich cat# P4850-5ML) was added, and cell lysate was incubated at 55 °C overnight. The following day, 5 µL RNase A (Sigma-Aldrich cat# R4642-50MG) was added, and the cell lysate was incubated at 37 °C for 30 min. Genomic DNA was precipitated by adding 4 µL Glycoblue (Fisher cat# AM9515), 40 µL 3M Sodium Acetate, and 1 mL ice-cold 100% ethanol. After incubating at −80 °C for 1 h, DNA was pelleted by centrifugation at 20,000 g for 30 min at 4 °C. The DNA pellet was washed twice with 70% ethanol, and then resuspended in 200 µL Buffer EB (Qiagen cat# 19086).

### RNA Purification

11-14 days post-transfection, cell pellets containing 1×10^6^ cells each were resuspended in 350 µL Trizol solution (Fisher cat# 15596026) and stored at −80 °C until RNA extraction. Frozen samples were allowed to warm to room temperature, and then 350 µL cell solution was transferred to a Phase-Lock gel tube (Fisher cat# NC1093153). 70 µL chloroform was added to each Phase-lock tube and shaken vigorously, followed by a 2 min incubation at room temperature. Tubes were centrifuged at 12,000 x g for 10 min at 4 °C. Following centrifugation, the aqueous phase was decanted from each tube and transferred to a new tube. 350 µL 70% ethanol was added and mixed well, and the solution was transferred to a Qiagen RNeasy-mini spin column. Samples were centrifuged at 13,000 x g for 15 s, and the flow-through was discarded. 350 µL Buffer RW1 was added to each column, samples were centrifuged at 13,000 x g for 15 s, and the flow-through was discarded. This Buffer RW1 wash was repeated once more for a total of 2 washes. Next, 500 µL Buffer RPE was added to each column, samples were centrifuged at 13,000 x g for 15 s, and the flow-through was discarded. This Buffer RPE wash was repeated once more for a total of 2 washes. After the last RPE wash, the column was centrifuged for an additional 2 min at 13,000 x g to remove residual ethanol. Samples were eluted in 40 µL RNase-free H_2_O.

To ensure that the RNA preparation was DNA-free, we utilized the Ambion TURBO DNA-free kit protocol (Thermo Fisher Scientific cat# AM1907). Following DNase treatment, RNA was transferred to a fresh tube, and the concentration was quantified on the Nanodrop.

### Amplicon Library Preparation

For DNA libraries, unique Molecular Identifiers (UMIs) and the inner portion of the P5 sequencing adapter were added. 20 µg of genomic DNA was digested with PstI (NEB cat# R3140L) for 1 h at 37 °C, and then purified using the Zymo Clean and Concentrate-25 (Zymo Research cat# D4034) protocol. One cycle of PCR was performed with the following conditions: 8 replicate 50 µL reactions were prepared, each containing 500 ng PstI digested DNA, 1x Phusion Hot Start Flex Mastermix (NEB cat# M0536L), and 200 nM of the primer P5_Plasmid_8N/9N/10N, and incubated at 98 °C for 5 min, 60 °C for 1 min, and 72 °C for 10 min. Replicate reactions were combined and then purified using the Zymo Clean and Concentrate-5 (Zymo Research cat# D4014) protocol, eluting the DNA in 20 µL.

For RNA libraries, cDNA was synthesized using the Superscript IV First Strand Synthesis kit (Invitrogen), with 5 µg RNA template and 2 µM primer P5_barcode_0N/1N/2N (containing a truncated sequencing adapter) in two replicate reactions per sample. RNA was first incubated with primers and dNTPs at 60 °C for 10 min, then placed on ice for 1 min. The remaining RT reagents were added, and samples were incubated at 55 °C for 10 min, 80 °C for 10 min, and then cooled to 4 °C. Next, 1 µL RNaseH was added to each reaction, and incubated at 37 °C for 20 min. Singlestranded cDNA was purified using the Zymo Clean and Concentrate-5 protocol (Zymo Research cat# D4014), using 7 volumes of DNA binding buffer and eluting in 10 µL Zymo DNA elution buffer. Unique Molecular Identifiers (UMIs) and the inner portion of the P7 sequencing adapter were added to each single-stranded cDNA molecule using 1 cycle of PCR with the following conditions: 2 replicate 50 µL reactions were prepared, each containing 5 µL cDNA, 1x Phusion Hot Start Flex Mastermix (NEB cat# M0536L), and 200 nM of the primer P7_Plasmid_8N/9N/10N, and incubated at 98 °C for 5 min, 64 °C for 5 min, and 72 °C for 5 min. Replicate reactions were combined and then purified using the Zymo Clean and Concentrate-5 (Zymo Research cat# D4014) protocol, eluting the DNA in 20 µL.

For inverse PCR (iPCR) libraries, 40 µg genomic DNA was digested with DpnII for 2 h at 37 °C. Digested DNA was purified using the Zymo Clean & Concentrate-25 column protocol, and digestion was verified by running 100 ng DpnII digested DNA out on a 1% agarose gel. Intramolecular DpnII ligation was performed using DpnII digested DNA at a concentration of 5 µg/mL, and T4 DNA ligase at a concentration of 10,000 U/mL. Ligation reactions were incubated overnight at 4 °C, and ligation products were purified using the Zymo Clean & Concentrate-25 column protocol.

DNA, RNA, and iPCR libraries then amplified using a nested PCR approach to add full Illumina sequencing adapters in two stages. To add the inner P5 and P7 sequencing adapters (DNA and RNA samples already had P5 or P7 added, respectively), samples were amplified for 20-30 PCR cycles. 8 replicate 50 µL reactions were prepared, each containing 2 µL DNA, 1x Phusion Hot Start Flex Mastermix (NEB cat# M0536L), 200 nM of the appropriate P5 and P7 primers for each library type (**Supplemental Table S5**), and incubated 1 cycle at 98 °C for 5 min; 20-30 cycles (sample dependent) at 98 °C for 15 s, 55 °C for 15 s, and 72 °C for 30 s; and 1 cycle of 72 °C for 10 min. Replicate reactions were combined and then purified using the Zymo Clean and Concentrate-5 (Zymo Research cat# D4014) protocol, eluting the DNA in 20 µL.

The remaining (outer) adapter sequences with indexing barcodes were added to each library using 10 cycles of PCR with the following conditions: one 50 µL reaction was prepared per library, each containing 1 µL DNA purified from the previous round of PCR, 1x Phusion Hot Start Flex Mastermix (NEB cat# M0536L), 200 nM of each indexed P5 and P7 primers (e.g. P5_amplicon_S502 and P7_Amplicon_N704, **Supplemental Table S5**), and incubated 1 cycle at 98 °C for 5 min, 10 cycles at 98 °C for 15 s, 71 °C for 15 s, and 72 °C for 30 s, and 1 cycle of 72 °C for 10 min. Final DNA libraries were purified using the Zymo Clean and Concentrate-5 (Zymo Research cat# D4014) protocol, eluting the DNA in 20 µL. Completed libraries were quantified using the Qubit dsDNA HS (Fisher cat# Q32851) kit protocol.

### Single Cell RNA-seq (scRNA-seq) Library Preparation

Cells were transfected as described above (**Supplemental Table S1**). To enrich for cells that received the transposase construct, mKate-positive cells were sorted into new plates 24 h after transfection using a Sony SH800 cell sorter as described above, except that the 665±30 nm optical filter was used to gate for mKate fluorescence, and expanded.

scRNA-seq expression libraries were generated using the 10x Chromium NextGem Single Cell 3’ workflow (10x Genomics cat# 1000128). For experiment 4, an additional sort for GFP-positive cells was performed on day 4, and 5700 cells were collected for scRNA-seq (**Supplemental Table S4**) while the remaining cells were expanded in culture for an additional 7 d, at which point cell pellets were collected for RNA, DNA, and iPCR libraries.

For experiment 5, cell pellets were collected for RNA, DNA, and iPCR libraries and cells were frozen 14 days post-transfection. After thawing cells and expanding for 2 days, cells were collected from each of 4 separate transfections, and pooled for scRNA-seq libraries (**Supplemental Table S4**). Additional pellets were collected post-thaw for additional RNA, DNA, and iPCR libraries.

Separate libraries enriched for reporter transcripts were generated from the cDNA produced in the Post GEM–RT Cleanup & cDNA Amplification step of the 10x Chromium Single Cell 3’ (v3.1) library protocol using PCR with the following conditions: 8 replicate 25 µL reactions were prepared, each containing 1 µL amplified 10x Chromium Single Cell cDNA, 1x Phusion Hot Start Flex Mastermix (NEB cat# M0536L), 500 nM of the primer P5_Halfsite, and 500 nM of the primer P7_10xSBbarcodeV2_0N, and incubated 1 cycle at 98 °C for 5 min; 8 cycles (sample dependent) at 98 °C for 15 s, 66 °C for 15 s, and 72 °C for 30 s; and 1 cycle of 72 °C for 10 min. Replicate reactions were combined and then purified using the Zymo Clean and Concentrate-5 (Zymo Research cat# D4014) protocol, eluting the DNA in 20 µL. Completed indexed adapter sequences were added to each library during a final 10 cycles of PCR using the conditions described in DNA Library Preparation above. Completed libraries were quantified using the Qubit dsDNA HS (Fisher cat# Q32851) kit protocol.

### Sequencing and analysis

Illumina libraries were generated and sequenced on an Illumina NextSeq 500. Reads were demultiplexed by a standard pipeline using Illumina bcl2fastq v2.20 requiring a perfect match to indexing BC sequences.

DNA, RNA, iPCR, and enriched scRNA-seq libraries were processed by a custom pipeline (**Supplemental Table S2**). Read pairs whose sequence comprised >75% G bases were dropped. PCR primer sequence was removed and UMI and cell barcodes (cell BC) were extracted using UMI-tools v1.0.1 (Smith et al. 2017) including the option ’--quality-filter-threshold=30’. Reporter barcodes (BCs) were extracted from the expected position from read pairs matching the expected template sequence with fewer than 10% mismatched bases. BCs were required to have 2 bases or fewer with base quality score below 30. Reporter BCs, cell BCs, and UMIs were each deduplicated using a directed adjacency approach based on that of UMI-tools (Smith et al. 2017).

iPCR libraries were trimmed to remove plasmid sequences, including the potential for digestion at a secondary DpnII site using cutadapt v2.9 (Martin 2011). Reads were then mapped to a hg38 reference genome augmented with transposon and sleeping beauty sequences using BWA v.0.7.12 (Li and Durbin 2009). Libraries where read 1 was sequenced to 24 bp or more beyond the end of the plasmid sequence were mapped in paired-end mode using BWA-MEM with -Y option. Otherwise read 2 was mapped in single-end mode using BWA aln and samse. Reads without reporter BCs, aligned with insertions or deletions, with >10% mismatch rate, with mapping quality <10, or with >1 kb between mates (for paired mapping) were excluded from further analysis. The integration insertion site was defined as the 5’ mapping site of read 2. Reporter BCs were additionally deduplicated grouped by coordinates. Sites with the same reporter BC within ±5 bp of the same BC were collapsed, integrations of different BCs within ±5 bp were excluded. Integrations with <2 reads, representing <1% of the total coverage at a given genomic position, or BCs found at multiple sites were excluded.

Read counts for DNA, RNA, and iPCR libraries were normalized per 1M sequenced reads and merged based on reporter BC. Missing RNA counts were imputed as 0, and only BCs with >10 DNA reads and an integration site were considered. Reporter activity was computed as log_2_(RNA/DNA + 1).

DNase-seq data (K562-DS9764) for K562 was downloaded from https://www.encodeproject.org and processed using a standard pipeline (https://github.com/mauranolab/mapping/tree/master/dnase). DNase I hypersensitive sites were identified using hotspot v1 (John et al. 2011) hotspot peaks (1% FDR). CTCF binding sites determined by ChIP-seq were taken from previously published work (Maurano et al. 2015).

### Clonal inference

Cells and reporter BCs deriving from a single initial transfected clone were derived from the enriched scRNA-seq libraries. First, we constructed a bipartite graph whose nodes were cell BCs and reporter BCs, connected by edges weighted by the pair’s UMI count. Edges with <2 UMI were dropped. The Jaccard index of reporter BC overlap for all pairs of cells within a clone was computed as the sum of edge weights connecting the cells to shared BCs divided by the sum of the UMIs for both cells. For all cell pairs whose Jaccard index was below 30%, the edge with the lowest weight between either cells was removed for each shared BC.

For Experiment 5, where scRNA-seq data was generated from a superloaded pool of 4 independent transfections, each reporter BC was labelled by its known transfection based on the union of all DNA/RNA/iPCR data. BCs found in more than one transfection were removed from the graph. Edges connecting a cell BC to a reporter BC from a transfection representing <80% of the cell’s total UMI were trimmed. Nodes directly connected to two different transfections (i.e. doublets or reporter BC collisions) were dropped.

To reduce the impact of chimeric PCR artifacts, edges representing <5% of total UMI for a given reporter BC or <2% for a given cellBC. Reporter BCs mapping to multiple integration sites or found in multiple transfections were removed. Finally, edges bridging two independent communities of 2 or more nodes with <20% centrality and representing <10% of each community’s UMIs were pruned. Unconnected nodes were pruned. The remaining connected communities were defined as clones.

### scRNA-seq analysis

scRNA-seq 3’ libraries were analyzed using Cell Ranger v.4.0.0 (Zheng et al. 2017). A reference was constructed against hg38 and transposon sequences as above using. Ensembl release 93 was used for gene annotations. Only cellBCs contained in the whitelist of non-empty cellBCs and absent from blacklists of poor quality cellBCs with few UMIs and/or too many pSB reads were considered on further analysis.

### Reporter effects on endogenous gene expression

We explored the reporter impact on genes whose TSS lay within 250 kb from a reporter. For each reporter and gene, we compared the expression on the set of perturbed cells (those belonging to the reporter’s clone) against all other cells. Only genes with Ensembl category of protein_coding or lincRNA were considered. Only cells included in both single-cell expression and clone assignments were used. To avoid potential confounding from nearby reporter insertions in the same clone, we discarded any reporters with a second reporter within 500 kb. Finally, tests with <3 perturbed cells, average target gene expression in the perturbed or unperturbed cells of <10 UMI, or overall average expression <0.05 UMI were excluded from the analysis.

Differential Expression of Clonal Alterations Local effects (DECAL) models the reporter effect using a negative binomial (or Gamma-Poisson) regression with regularized dispersion estimate:

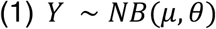

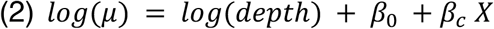

Where Y are the observed counts for a particular gene across all cells, µ is the expected average gene UMIs, θ is the gene UMI distribution dispersion, *β*_0_ and *β*_*c*_ are the regression coefficients, *depth* is each cell’s UMIs, and *X* is an indicator vector that is 1 if the cell belongs to the reporter clone (perturbed) or 0 if it does not (unperturbed).

In order to estimate the distribution dispersion (θ) of each gene, we employed the approach of Hafemeister & Satija (Hafemeister and Satija 2019) of fitting a Poisson regression 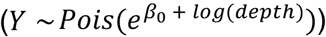 for a random subset of 2000 genes and estimating θ using maximum likelihood. Then, we expanded the estimation to all genes with average expression ≥0.05 UMIs by fitting a kernel regression of θ in relation to the gene average expression.

Perturbation (*β*_*c*_) significance was estimated by two-tailed p-value based on Student’s t-distribution. Storey’s q-value approach was used for multiple testing correction (Storey and Tibshirani 2003) for each transfection individually. Tests with q-value <0.05 were considered significant.

We performed simulations to estimate detection power over a range of clone sizes (n) and effect sizes using equation 1 above with a fixed dispersion (θ) of 100. For each simulation condition, we generated UMI counts for 1000 genes, and the same number of cells as the actual scRNA-seq dataset. Expected UMIs (µ) ranged from the minimum and maximum observed values in the actual scRNA-seq dataset in a logarithm scale. For each gene, n cells were sampled and their UMI counts altered by the defined effect size. The resulting simulations were evaluated by our analysis algorithm given only the simulated count matrix and cells assignment as perturbed and unperturbed.

Contact matrices were taken from (Rao et al. 2014) GSE63525_K562_intrachromosomal_contact_matrices.tar.gz), KR normalized, and Armatus v2.2 (Filippova et al. 2014) was used to identify TADs with gamma=0.5.0 and a resolution of 5 kb. Results were lifted over to hg38 using liftOver.

## Supporting information

SOM

## Software availability

Code used in sequencing data processing is available at GitHub (https://github.com/mauranolab/mapping/tree/master/transposon). Code used for scRNA-seq differential expression analysis is available at GitHub (https://github.com/mauranolab/decal).

## Data access

All raw and processed sequencing data generated in this study have been submitted to the NCBI Gene Expression Omnibus (GEO; https://www.ncbi.nlm.nih.gov/geo/) under accession number GSE179485.

## Acknowledgements

We thank Ran Brosh for careful reading of the manuscript and assistance analyzing flow cytometry data. This work was partially funded by NIH grant R35GM119703 to M.T.M..

## References

Akhtar W, de Jong J, Pindyurin AV, Pagie L, Meuleman W, de Ridder J, Berns A, Wessels LFA, van Lohuizen M, van Steensel B. 2013. Chromatin position effects assayed by thousands of reporters integrated in parallel. Cell 154: 914–927.

Bell AC, Felsenfeld G. 2000. Methylation of a CTCF-dependent boundary controls imprinted expression of the Igf2 gene. Nature 405: 482–485.

Biddy BA, Kong W, Kamimoto K, Guo C, Waye SE, Sun T, Morris SA. 2018. Single-cell mapping of lineage and identity in direct reprogramming. Nature 564: 219–224.

Chung JH, Whiteley M, Felsenfeld G. 1993. A 5’ element of the chicken beta-globin domain serves as an insulator in human erythroid cells and protects against position effect in Drosophila. Cell 74: 505–514.

de Wit E, Vos ESM, Holwerda SJB, Valdes-Quezada C, Verstegen MJAM, Teunissen H, Splinter E, Wijchers PJ, Krijger PHL, de Laat W. 2015. CTCF Binding Polarity Determines Chromatin Looping. Molecular Cell 60: 676–684.

Dickson J, Gowher H, Strogantsev R, Gaszner M, Hair A, Felsenfeld G, West AG. 2010. VEZF1 elements mediate protection from DNA methylation. PLoS Genetics 6: e1000804.

Dixon JR, Selvaraj S, Yue F, Kim A, Li Y, Shen Y, Hu M, Liu JS, Ren B. 2012. Topological domains in mammalian genomes identified by analysis of chromatin interactions. Nature 485: 376–380.

Filippova D, Patro R, Duggal G, Kingsford C. 2014. Identification of alternative topological domains in chromatin. Algorithms Mol Biol 9: 14.

Gasperini M, Hill AJ, McFaline-Figueroa JL, Martin B, Kim S, Zhang MD, Jackson D, Leith A, Schreiber J, Noble WS, et al. 2019. A Genome-wide Framework for Mapping Gene Regulation via Cellular Genetic Screens. Cell 176: 1516.

Guo Y, Xu Q, Canzio D, Shou J, Li J, Gorkin DU, Jung I, Wu H, Zhai Y, Tang Y, et al. 2015. CRISPR Inversion of CTCF Sites Alters Genome Topology and Enhancer/Promoter Function. Cell 162: 900–910.

Hafemeister C, Satija R. 2019. Normalization and variance stabilization of single-cell RNA-seq data using regularized negative binomial regression. Genome Biol 20: 296.

Halow JM, Byron R, Hogan MS, Ordoñez R, Groudine M, Bender MA, Stamatoyannopoulos JA, Maurano MT. 2021. Tissue context determines the penetrance of regulatory DNA variation. Nat Commun 12: 2850.

Herranz D, Ambesi-Impiombato A, Palomero T, Schnell SA, Belver L, Wendorff AA, Xu L, Castillo-Martin M, Llobet-Navás D, Cordon-Cardo C, et al. 2014. A NOTCH1-driven MYC enhancer promotes T cell development, transformation and acute lymphoblastic leukemia. Nat Med 20: 1130–1137.

Huang H, Zhu Q, Jussila A, Han Y, Bintu B, Kern C, Conte M, Zhang Y, Bianco S, Chiariello AM, et al. 2021. CTCF mediates dosage-and sequence-context-dependent transcriptional insulation by forming local chromatin domains. Nat Genet.

Jee J, Rasouly A, Shamovsky I, Akivis Y, Steinman SR, Mishra B, Nudler E. 2016. Rates and mechanisms of bacterial mutagenesis from maximum-depth sequencing. Nature 534: 693–696.

John S, Sabo PJ, Thurman RE, Sung M-H, Biddie SC, Johnson TA, Hager GL, Stamatoyannopoulos JA. 2011. Chromatin accessibility pre-determines glucocorticoid receptor binding patterns. Nature Genetics 43: 264–268.

Li CL, Xiong D, Stamatoyannopoulos G, Emery DW. 2009. Genomic and functional assays demonstrate reduced gammaretroviral vector genotoxicity associated with use of the cHS4 chromatin insulator. Mol Ther 17: 716–724.

Li H, Durbin R. 2009. Fast and accurate short read alignment with Burrows-Wheeler transform. Bioinformatics 25: 1754–1760.

Liu M, Maurano MT, Wang H, Qi H, Song C-Z, Navas PA, Emery DW, Stamatoyannopoulos JA, Stamatoyannopoulos G. 2015. Genomic discovery of potent chromatin insulators for human gene therapy. Nature Biotechnology 33: 198–203.

Lu R, Neff NF, Quake SR, Weissman IL. 2011. Tracking single hematopoietic stem cells in vivo using high-throughput sequencing in conjunction with viral genetic barcoding. Nat Biotechnol 29: 928–933.

Maricque BB, Chaudhari HG, Cohen BA. 2018. A massively parallel reporter assay dissects the influence of chromatin structure on cis-regulatory activity. Nat Biotechnol.

Martin M. 2011. Cutadapt removes adapter sequences from high-throughput sequencing reads. EMBnet.journal 17: 10–12.

Mátés L, Chuah MKL, Belay E, Jerchow B, Manoj N, Acosta-Sanchez A, Grzela DP, Schmitt A, Becker K, Matrai J, et al. 2009. Molecular evolution of a novel hyperactive Sleeping Beauty transposase enables robust stable gene transfer in vertebrates. Nature Genetics 41: 753–761.

Maurano MT, Wang H, John S, Shafer A, Canfield T, Lee K, Stamatoyannopoulos JA. 2015. Role of DNA Methylation in Modulating Transcription Factor Occupancy. Cell reports 12: 1184–1195.

Moudgil A, Wilkinson MN, Chen X, He J, Cammack AJ, Vasek MJ, Lagunas T, Qi Z, Lalli MA, Guo C, et al. 2020. Self-Reporting Transposons Enable Simultaneous Readout of Gene Expression and Transcription Factor Binding in Single Cells. Cell 182: 992-1008.e21.

Phillips JE, Corces VG. 2009. CTCF: master weaver of the genome. Cell 137: 1194–1211.

Qi H, Liu M, Emery DW, Stamatoyannopoulos G. 2015. Functional validation of a constitutive autonomous silencer element. PLoS One 10: e0124588.

Radtke I, Mullighan CG, Ishii M, Su X, Cheng J, Ma J, Ganti R, Cai Z, Goorha S, Pounds SB, et al. 2009. Genomic analysis reveals few genetic alterations in pediatric acute myeloid leukemia. Proc Natl Acad Sci U S A 106: 12944–12949.

Rao SSP, Huntley MH, Durand NC, Stamenova EK, Bochkov ID, Robinson JT, Sanborn AL, Machol I, Omer AD, Lander ES, et al. 2014. A 3D map of the human genome at kilobase resolution reveals principles of chromatin looping. Cell 159: 1665–1680.

Smith T, Heger A, Sudbery I. 2017. UMI-tools: modeling sequencing errors in Unique Molecular Identifiers to improve quantification accuracy. Genome Res 27: 491–499.

Storey JD, Tibshirani R. 2003. Statistical significance for genomewide studies. Proceedings of the National Academy of Sciences of the United States of America 100: 9440–9445.

Sur IK, Hallikas O, Vähärautio A, Yan J, Turunen M, Enge M, Taipale M, Karhu A, Aaltonen LA, Taipale J. 2012. Mice lacking a Myc enhancer that includes human SNP rs6983267 are resistant to intestinal tumors. Science 338: 1360–1363.

Walters MC, Fiering S, Bouhassira EE, Scalzo D, Goeke S, Magis W, Garrick D, Whitelaw E, Martin DI. 1999. The chicken beta-globin 5’HS4 boundary element blocks enhancer-mediated suppression of silencing. Mol Cell Biol 19: 3714–3726.

Walters MC, Magis W, Fiering S, Eidemiller J, Scalzo D, Groudine M, Martin DI. 1996. Transcriptional enhancers act in cis to suppress position-effect variegation. Genes Dev 10: 185–195.

Weinreb C, Rodriguez-Fraticelli A, Camargo FD, Klein AM. 2020. Lineage tracing on transcriptional landscapes links state to fate during differentiation. Science 367: eaaw3381.

Weintraub H. 1988. Formation of stable transcription complexes as assayed by analysis of individual templates. Proceedings of the National Academy of Sciences of the United States of America 85: 5819–5823.

West AG, Gaszner M, Felsenfeld G. 2002. Insulators: many functions, many mechanisms. Genes Dev 16: 271–288.

Zheng GXY, Terry JM, Belgrader P, Ryvkin P, Bent ZW, Wilson R, Ziraldo SB, Wheeler TD, McDermott GP, Zhu J, et al. 2017. Massively parallel digital transcriptional profiling of single cells. Nat Commun 8: 14049.

