## Supplementary material for "Genomic context sensitivity of insulator function": SOM

**Supplemental Material for Ribeiro-dos-Santos et al.**
**“Genomic context sensitivity of insulator function”**

### Supplemental Figures

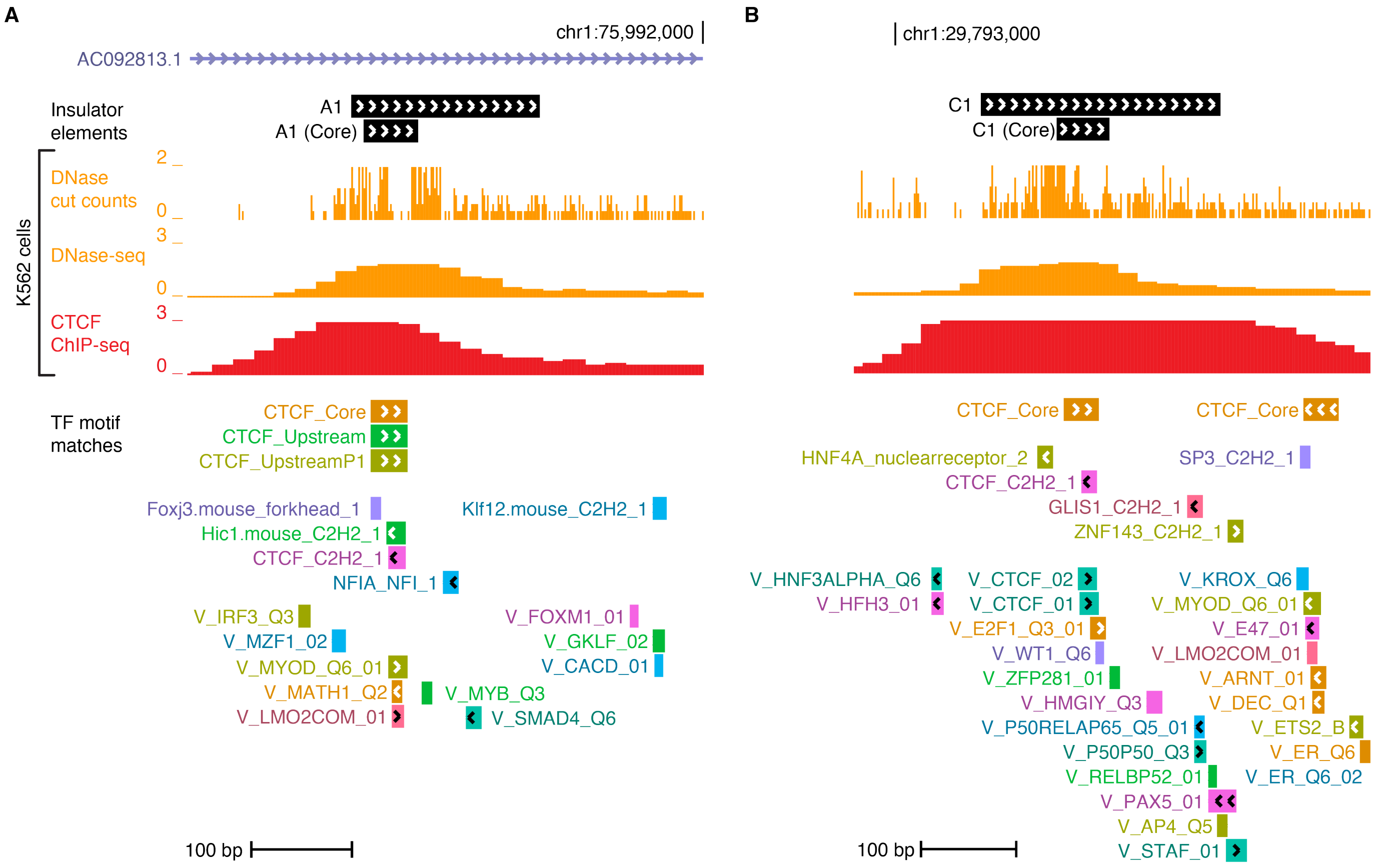

**Supplemental Fig. S1. A1 and C1 genomic insulator elements.**
(A-B) Shown are full A1 (A) and C1 (B) insulator elements from (Liu et al. 2015). A1 and C1 core elements have been truncated to just the CTCF footprint. Shown are DNase-seq cut counts, DNase-seq density, and CTCF ChIP-seq tracks from K562 cells. Below are TF motif matches using FIMO ($P < 10^{-5}$).

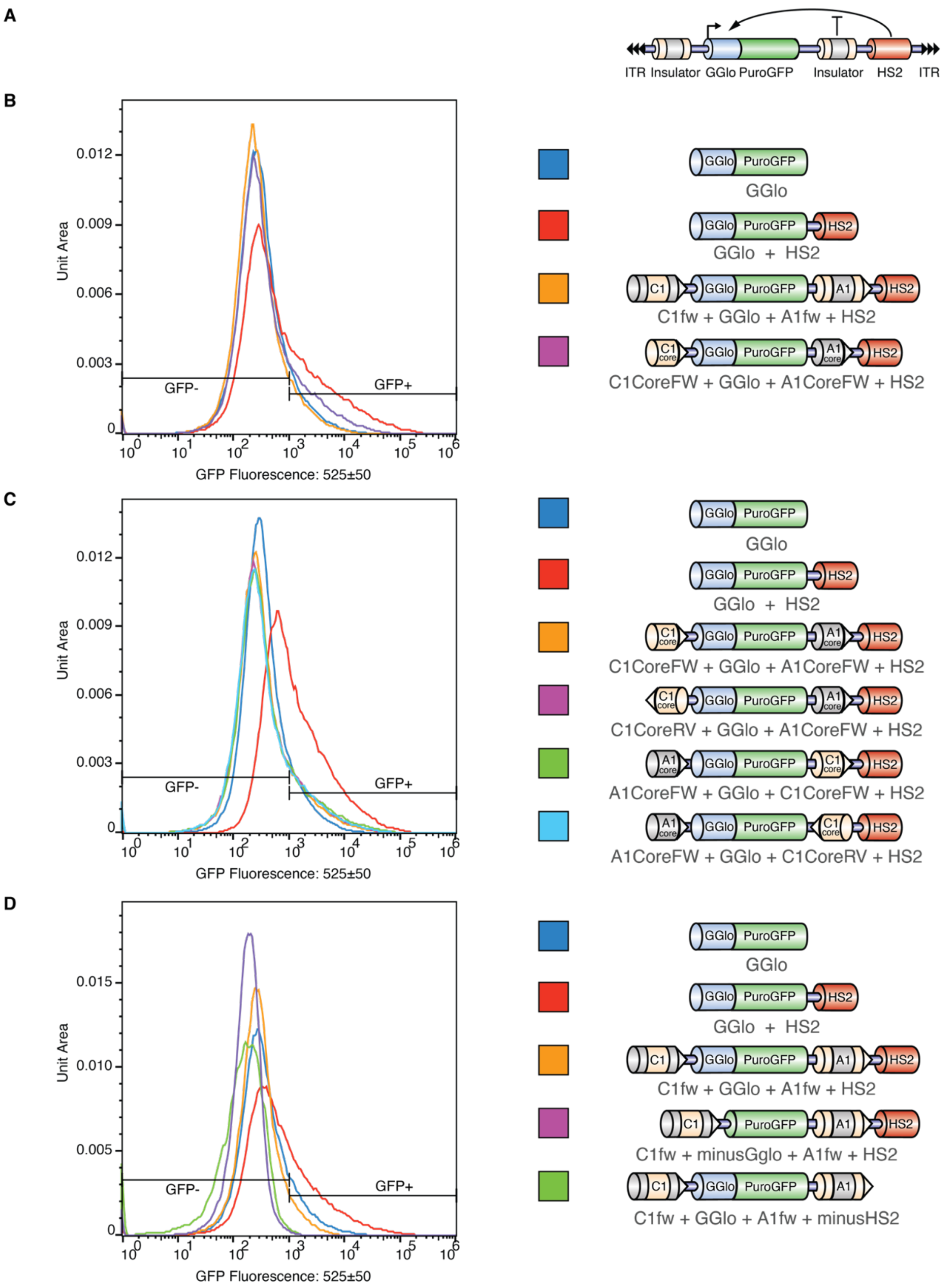

**Supplemental Fig. S2. Characterization of enhancer blocker reporter using flow cytometry.**
(A) Reporter scheme consisting of ɣ-globin promoter driving PuroGFP expression. An intervening CTCF site between the promoter and HS2 acts as an enhancer blocker. ITR, Sleeping Beauty inverted terminal repeats; GGlo, ɣ-globin promoter; HS2, ß-globin hypersensitive site 2 enhancer. (B-D) GFP positive cells were measured by flow cytometry. GGlo and GGlo+HS2 were repeated in all experiments. (B) The A1 insulator behaves as a strong enhancer blocker element. Truncating A1 to just the CTCF site (A1core, see **Supplemental Fig. S1**) retains potent enhancer blocker activity. (C) Enhancer blocker activity is independent of the orientation of the insulator element. C1core (**Supplemental Fig. S1**) is a strong enhancer blocker element. (D) Reporters without GGlo or HS2 each demonstrate no GFP expression.

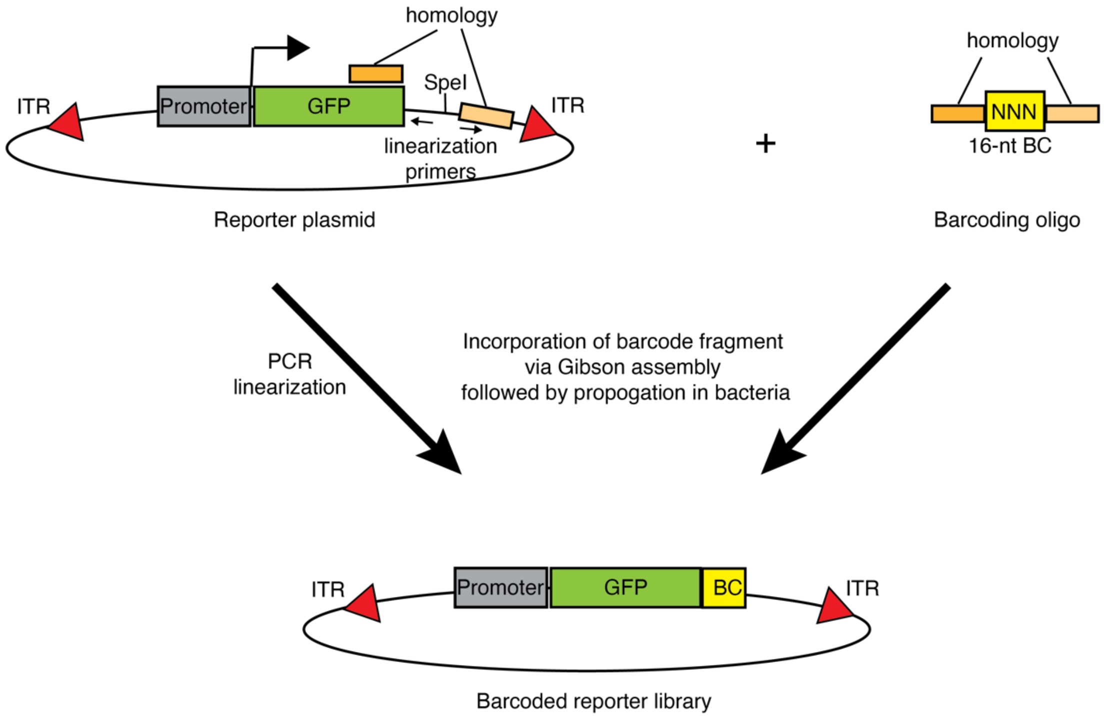

**Supplemental Fig. S3. Reporter plasmid barcoding strategy.**
Libraries of barcoded reporter plasmids were generated by PCR linearization followed by incorporation of synthetic oligonucleotide containing a 16-nt random reporter BC sequence using Gibson assembly. Orange rectangles indicate homology regions for reporter BC cloning. ITR, Sleeping Beauty inverted terminal repeats.

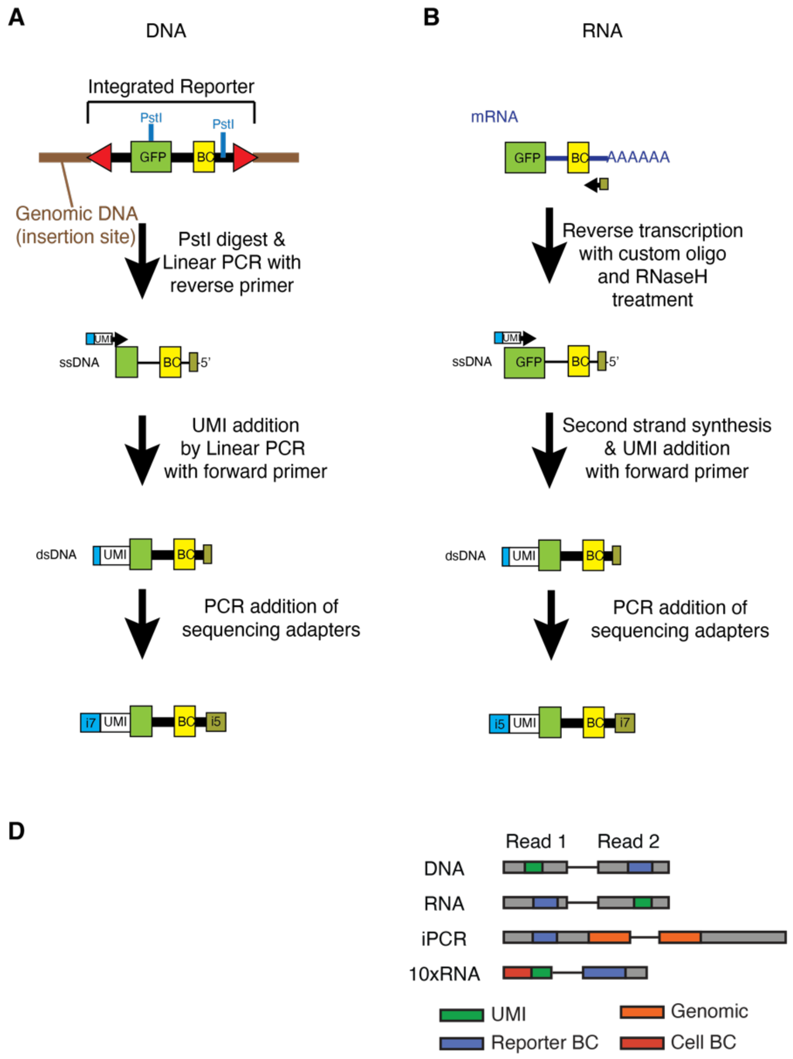

**Supplemental Fig. S4. Amplicon library construction approach.**
(A-C) Preparation of Illumina sequencing libraries to quantify barcoded reporter representation from DNA, expression from RNA, and integration location using inverse PCR (iPCR). DNA libraries utilized a PstI digest to limit template size, followed by a single linear amplification to add UMIs, and finally an Exo I digest to prevent any amplification from untemplated primers. RNA libraries incorporated UMIs during second strand synthesis. (D) Schematic of unique molecular identifier (UMI, DNA, RNA, and 10x scRNA-seq libraries), Reporter BC, Cell BC (10x scRNA-seq libraries), and genomic sequence (iPCR libraries). The number of N nucleotides added to each individual sample varied between 8-12 nt (DNA and RNA) or 0-2 nt (iPCR) to increase diversity on the sequencing flow cell. Not shown: for DNA libraries, there can be an additional 0-2 nt of UMI on R2.

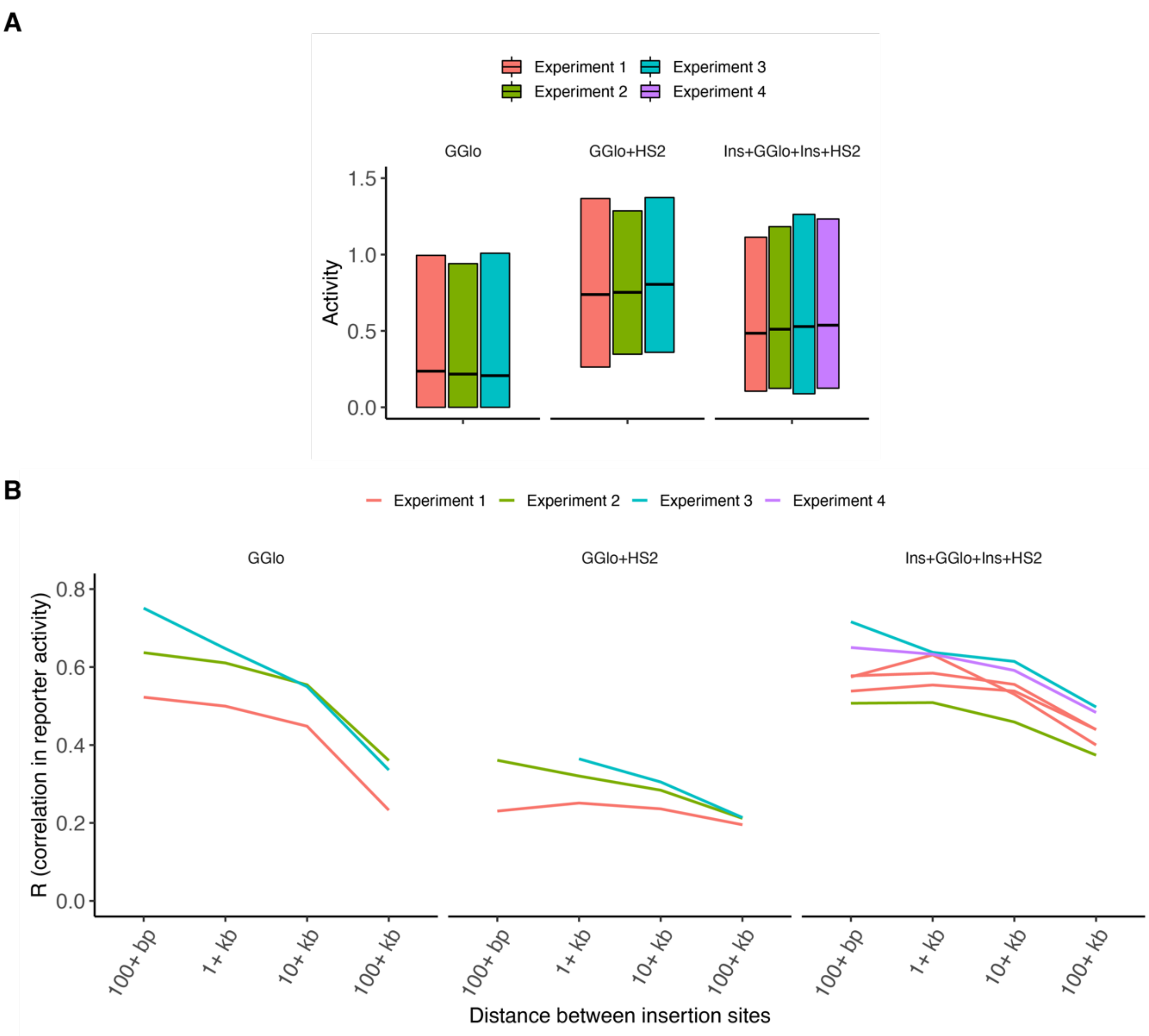

**Supplemental Fig. S5. Integrated barcoded reporter assay.**
(A) Average activity by reporter class and experiment. (B) Correlation in activity for nearby insertions by reporter and experiment. Bins with fewer than 100 datapoints were omitted.

**A**

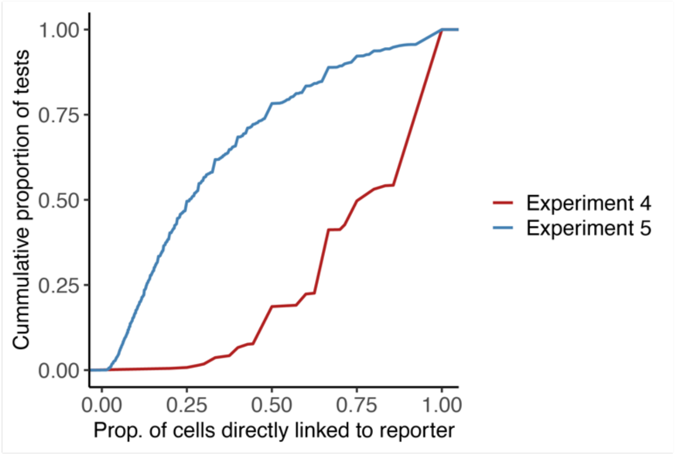

**B**

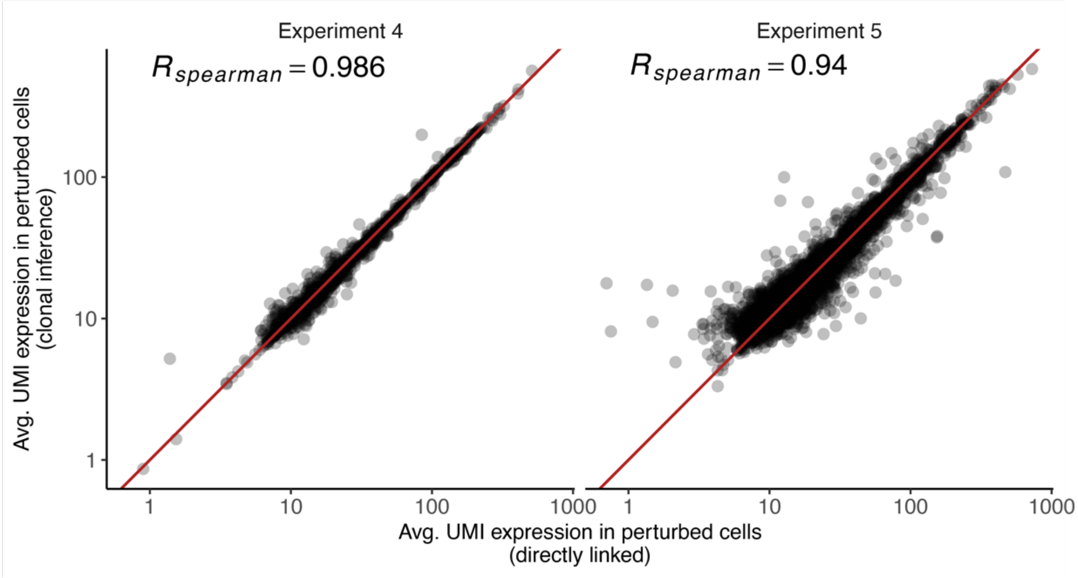

**Supplemental Fig. S6. Clonal inference.**

(A) Cumulative plot showing the proportion of cells with direct evidence for that reporter BC among all tests for differential expression. (B) Estimated expression of perturbed condition using cells where a reporter BC was directly measured (x-axis) vs. all cells in the same clonal lineage (y-axis). Tests with fewer than 20,000 UMIs on average among directly linked cells or fewer than 2 directly linked cells were omitted.

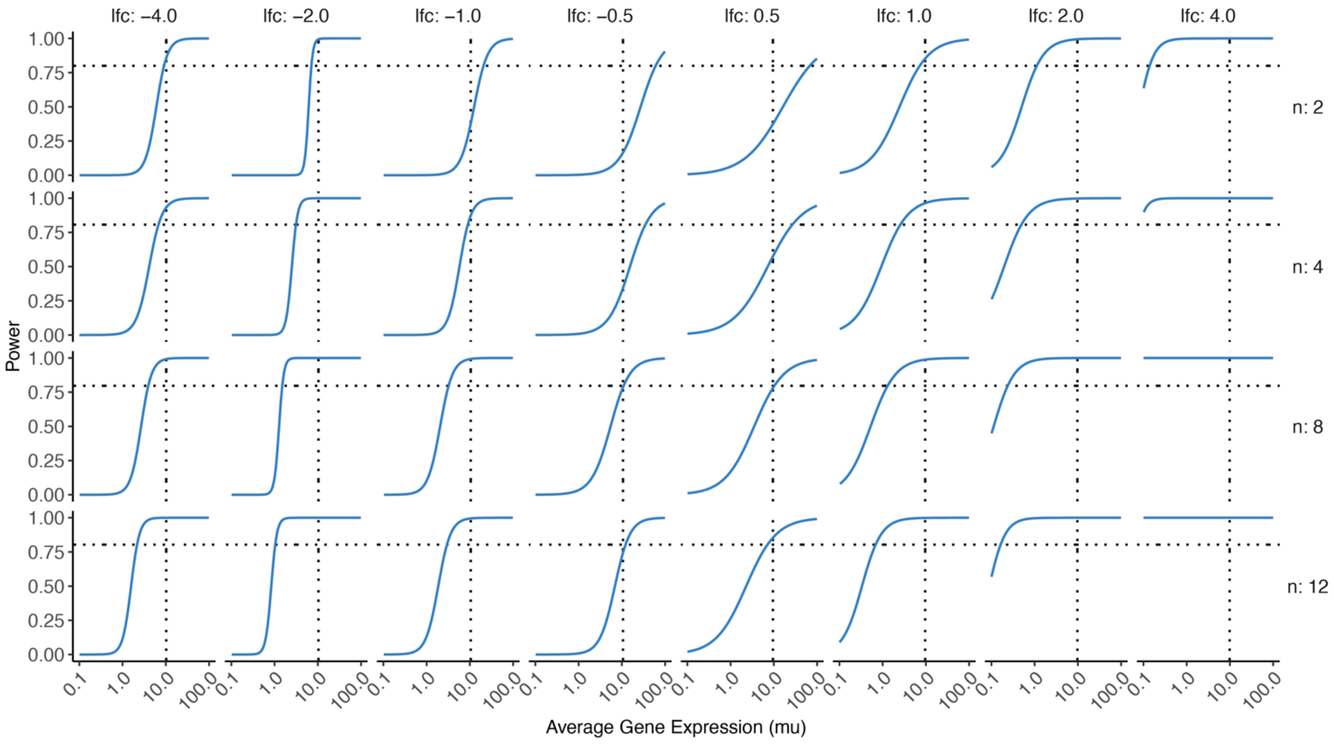

**Supplemental Fig. S7. Reporter impact on gene expression.**

(A) Detection power for different circumstances stratified by fold change of the perturbation (columns), number of cells in the clone (rows), and average expression (x-axis). Y-axis shows proportion of significant tests (q-value < 0.05) for each condition. Plotted is a logistic regression fit to the observed response. Horizontal dashed line shows 80% power, and vertical dashed line shows average gene expression of 10 UMIs.

### Supplemental Tables

#### Supplemental Table S1. Transfection summaries.

Summary of transfection and culture conditions. The Reporter Construct column summarizes the reporter configuration: GGlo, ɣ-globin promoter; HS2, ß-globin hypersensitive site 2 enhancer; A1fw, C1fw, A1rv, and C1rv, CTCF-binding insulator elements in the forward and reverse orientations (Liu et al. 2015); "Core" indicates the element has been truncated to just the CTCF recognition sequence (**Supplemental Fig. S1**). # Cells Seeded refers to the cells seeded on day 0 for Experiments 1-3, and the number of mKate+ cells on day 1 for Experiments 4-5. Puro indicates 2.5 $\mu$g/mL puromycin.

| Experiment | Plasmid # | Transfection | Reporter Construct | Amt. Transposon (pM) | Amt. Transposase (pM) | # Cells Seeded | Selection |
|---|---|---|---|---|---|---|---|
| GFP (Fig. 1) | pMH027 | | GGlo | 2,600 | 2,600 | 100,000 | puro from day 5 |
| GFP (Fig. 1) | pMH028 | | GGlo_HS2 | 2,600 | 2,600 | 100,000 | puro from day 5 |
| GFP (Fig. 1) | pMH032 | | C1fw_GGlo_A1fw_HS2 | 2,600 | 2,600 | 100,000 | puro from day 5 |
| GFP (Fig. 1) | pMH041 | | C1CoreFw_GGlo_A1CoreFw_HS2 | 2,600 | 2,600 | 100,000 | puro from day 5 |
| GFP (Fig. 1) | pMH027 | | GGlo | 2,600 | 2,600 | 100,000 | puro from day 5 |
| GFP (Fig. 1) | pMH028 | | GGlo_HS2 | 2,600 | 2,600 | 100,000 | puro from day 5 |
| GFP (Fig. 1) | pMH041 | | C1CoreFw_GGlo_A1CoreFw_HS2 | 2,600 | 2,600 | 100,000 | puro from day 5 |
| GFP (Fig. 1) | pMH043 | | A1CoreFw_GGlo_C1CoreFw_HS2 | 2,600 | 2,600 | 100,000 | puro from day 5 |
| GFP (Fig. 1) | pMH042 | | C1CoreRv_GGlo_A1CoreFw_HS2 | 2,600 | 2,600 | 100,000 | puro from day 5 |
| GFP (Fig. 1) | pMH044 | | A1CoreFw_GGlo_C1CoreRv_HS2 | 2,600 | 2,600 | 100,000 | puro from day 5 |
| GFP (Fig. 1) | pMH027 | | GGlo | 2,600 | 2,600 | 100,000 | puro from day 5 |
| GFP (Fig. 1) | pMH028 | | GGlo_HS2 | 2,600 | 2,600 | 100,000 | puro from day 5 |
| GFP (Fig. 1) | pMH032 | | C1fw_GGlo_A1fw_HS2 | 2,600 | 2,600 | 100,000 | puro from day 5 |
| GFP (Fig. 1) | pMH049 | | C1fw_GGlo_A1fw_minusHS2 | 2,600 | 2,600 | 100,000 | puro from day 5 |
| GFP (Fig. 1) | pMH050 | | C1fw_A1fw_HS2_minusGglo | 2,600 | 2,600 | 100,000 | puro from day 5 |
| Experiment 1 | pMH027 | | GGlo | 2,700 | 2,400 | 250,000 | puro from day 1 |
| Experiment 1 | pMH028 | | GGlo_HS2 | 2,200 | 2,400 | 250,000 | puro from day 1 |
| Experiment 1 | pMH032 | | C1fw_GGlo_A1fw_HS2 | 2,000 | 2,400 | 250,000 | puro from day 1 |
| Experiment 1 | pMH034 | | C1fw_GGlo_A1rv_HS2 | 2,000 | 2,400 | 250,000 | puro from day 1 |
| Experiment 2 | pMH027 | | GGlo | 1,800 | 2,400 | 90,000 | puro from day 1 |
| Experiment 2 | pMH028 | | GGlo_HS2 | 4,100 | 2,400 | 85,000 | puro from day 1 |
| Experiment 2 | pMH032 | | C1fw_GGlo_A1fw_HS2 | 2,200 | 2,400 | 190,000 | puro from day 1 |
| Experiment 2 | pMH034 | | C1fw_GGlo_A1rv_HS2 | 2,300 | 2,400 | 105,000 | puro from day 1 |
| Experiment 3 | pMH027 | | GGlo | 5,000 | 2,600 | 10,000 | puro from day 6 |
| Experiment 3 | pMH028 | | GGlo_HS2 | 5,000 | 2,600 | 10,000 | puro from day 6 |
| Experiment 3 | pMH032 | | C1fw_GGlo_A1rv_HS2 | 5,000 | 2,600 | 10,000 | puro from day 6 |
| Experiment 4 | pMH034 | | C1fw_GGlo_A1rv_HS2 | 5,000 | 2,600 | 10,000 | mKate sort day 1, GFP sort day 4 |
| Experiment 5 | pMH027 | A | GGlo_HS2 | 24,000 | 3,000 | 5,583 | mKate sort day 1 |
| Experiment 5 | pMH041 | B | C1CoreFw_GGlo_A1CoreFw_HS2 | 24,000 | 3,000 | 5,161 | mKate sort day 1 |
| Experiment 5 | pMH041 | C | C1CoreFw_GGlo_A1CoreFw_HS2 | 24,000 | 3,000 | 5,816 | mKate sort day 1 |
| Experiment 5 | pMH065 | D | C1CoreFw_GGlo_HS2_A1CoreFw | 24,000 | 3,000 | 4,045 | mKate sort day 1 |

**Supplemental Table S2. Sequencing libraries for DNA/RNA/iPCR/10xRNA experiments.**
Amplicon PCR libraries are listed by experiment. For Experiment 5, individual transfections are distinguished. Technical replicates (PCR library construction repeated on the same sample) are distinguished by letter suffix in "Sample #". Trim R1/R2 refers to the number of nt added to the R1 or R2 to increase sequencing diversity; UMI length indicates the portion of these that serve as a UMI (see **Methods**).

| Ex-peri-ment | Reporter Construct | Trans-fec-tion | Sample # | Type | Trim (R1/R2) | UMI length | Total read pairs (M) | Unique BC | Map-ping | Unique sites |
|---|---|---|---|---|---|---|---|---|---|---|
| 1 | GGlo | | BS02212A | DNA | 10/1 | 10 | 18.2 | 111,170 | NA | NA |
| 1 | GGlo | | BS02216A | DNA | 9/1 | 9 | 24.1 | 124,979 | NA | NA |
| 1 | GGlo | | BS02225A | iPCR | 0/1 | NA | 15.1 | 88,712 | SE | 50,586 |
| 1 | GGlo | | BS02225B | iPCR | 0/0 | NA | 6.8 | 82,983 | SE | 51,949 |
| 1 | GGlo | | BS02225C | iPCR | 0/0 | NA | 7.6 | 90,275 | SE | 59,441 |
| 1 | GGlo | | BS02225D | iPCR | 0/0 | NA | 8.1 | 91,620 | SE | 60,385 |
| 1 | GGlo | | BS02220A | RNA | 1/9 | 10 | 16.6 | 59,453 | NA | NA |
| 1 | GGlo | | BS02220B | RNA | 1/8 | 9 | 17.0 | 61,710 | NA | NA |
| 1 | GGlo_HS2 | | BS02213A | DNA | 9/2 | 9 | 14.1 | 84,216 | NA | NA |
| 1 | GGlo_HS2 | | BS02217A | DNA | 8/2 | 8 | 15.9 | 78,578 | NA | NA |
| 1 | GGlo_HS2 | | BS02226A | iPCR | 1/0 | NA | 17.6 | 32,889 | SE | 22,093 |
| 1 | GGlo_HS2 | | BS02226B | iPCR | 0/1 | NA | 7.4 | 31,720 | SE | 20,230 |
| 1 | GGlo_HS2 | | BS02226C | iPCR | 0/1 | NA | 7.1 | 34,301 | SE | 22,657 |
| 1 | GGlo_HS2 | | BS02221A | RNA | 0/10 | 10 | 13.6 | 73,488 | NA | NA |
| 1 | GGlo_HS2 | | BS02221B | RNA | 0/9 | 9 | 13.3 | 72,119 | NA | NA |
| 1 | C1fw_GGlo_A1fw_HS2 | | BS02214A | DNA | 8/0 | 8 | 14.9 | 45,307 | NA | NA |
| 1 | C1fw_GGlo_A1fw_HS2 | | BS02218A | DNA | 10/0 | 10 | 14.6 | 43,771 | NA | NA |
| 1 | C1fw_GGlo_A1fw_HS2 | | BS02227A | iPCR | 2/1 | NA | 22.1 | 27,836 | SE | 16,577 |
| 1 | C1fw_GGlo_A1fw_HS2 | | BS02227B | iPCR | 2/0 | NA | 11.3 | 26,440 | SE | 17,087 |
| 1 | C1fw_GGlo_A1fw_HS2 | | BS02227C | iPCR | 2/0 | NA | 11.3 | 24,073 | SE | 16,162 |
| 1 | C1fw_GGlo_A1fw_HS2 | | BS02222A | RNA | 0/8 | 8 | 11.4 | 26,929 | NA | NA |
| 1 | C1fw_GGlo_A1fw_HS2 | | BS02222B | RNA | 0/10 | 10 | 16.9 | 29,560 | NA | NA |
| 1 | C1fw_GGlo_A1rv_HS2 | | BS02215A | DNA | 10/0 | 10 | 13.8 | 64,818 | NA | NA |
| 1 | C1fw_GGlo_A1rv_HS2 | | BS02219A | DNA | 9/1 | 9 | 14.8 | 46,523 | NA | NA |
| 1 | C1fw_GGlo_A1rv_HS2 | | BS02228A | iPCR | 0/0 | NA | 19.5 | 42,792 | SE | 28,528 |
| 1 | C1fw_GGlo_A1rv_HS2 | | BS02228B | iPCR | 2/1 | NA | 11.1 | 39,048 | SE | 24,658 |
| 1 | C1fw_GGlo_A1rv_HS2 | | BS02228C | iPCR | 2/1 | NA | 7.8 | 32,706 | SE | 21,917 |
| 1 | C1fw_GGlo_A1rv_HS2 | | BS02223A | RNA | 2/9 | 11 | 17.5 | 56,186 | NA | NA |
| 1 | C1fw_GGlo_A1rv_HS2 | | BS02223B | RNA | 2/10 | 12 | 18.5 | 60,304 | NA | NA |
| 2 | GGlo | | BS02560A | DNA | 9/2 | 9 | 18.1 | 70,902 | NA | NA |
| 2 | GGlo | | BS02561A | DNA | 10/2 | 10 | 14.0 | 71,044 | NA | NA |
| 2 | GGlo | | BS02605A | iPCR | 0/0 | NA | 10.7 | 32,115 | SE | 15,117 |
| 2 | GGlo | | BS02605B | iPCR | 1/1 | NA | 6.8 | 24,168 | SE | 13,880 |
| 2 | GGlo | | BS02606A | iPCR | 2/0 | NA | 11.4 | 38,499 | SE | 20,250 |
| 2 | GGlo | | BS02539A | RNA | 2/8 | 10 | 4.9 | 34,904 | NA | NA |
| 2 | GGlo | | BS02539B | RNA | 2/8 | 10 | 5.6 | 42,237 | NA | NA |
| 2 | GGlo | | BS02540A | RNA | 1/8 | 9 | 5.0 | 30,078 | NA | NA |
| 2 | GGlo_HS2 | | BS02562A | DNA | 8/0 | 8 | 17.6 | 77,830 | NA | NA |
| 2 | GGlo_HS2 | | BS02563A | DNA | 9/0 | 9 | 15.6 | 77,992 | NA | NA |
| 2 | GGlo_HS2 | | BS02607A | iPCR | 0/1 | NA | 9.7 | 16,664 | SE | 7,488 |
| 2 | GGlo_HS2 | | BS02607C | iPCR | 0/1 | NA | 15.6 | 21,905 | SE | 10,303 |
| 2 | GGlo_HS2 | | BS02608A | iPCR | 1/0 | NA | 9.7 | 30,763 | SE | 18,109 |
| 2 | GGlo_HS2 | | BS02541A | RNA | 0/8 | 8 | 5.9 | 53,409 | NA | NA |
| 2 | GGlo_HS2 | | BS02542A | RNA | 2/9 | 11 | 4.5 | 53,612 | NA | NA |
| 2 | C1fw_GGlo_A1fw_HS2 | | BS02564A | DNA | 10/0 | 10 | 17.4 | 61,448 | NA | NA |
| 2 | C1fw_GGlo_A1fw_HS2 | | BS02565A | DNA | 8/1 | 8 | 18.3 | 56,228 | NA | NA |
| 2 | C1fw_GGlo_A1fw_HS2 | | BS02609A | iPCR | 2/1 | NA | 10.7 | 11,637 | SE | 5,282 |
| 2 | C1fw_GGlo_A1fw_HS2 | | BS02609B | iPCR | 0/0 | NA | 6.3 | 12,080 | SE | 5,966 |
| 2 | C1fw_GGlo_A1fw_HS2 | | BS02610A | iPCR | 1/1 | NA | 10.5 | 18,061 | SE | 9,893 |
| 2 | C1fw_GGlo_A1fw_HS2 | | BS02543A | RNA | 1/9 | 10 | 3.4 | 19,355 | NA | NA |
| 2 | C1fw_GGlo_A1fw_HS2 | | BS02543B | RNA | 1/9 | 10 | 4.4 | 26,838 | NA | NA |
| 2 | C1fw_GGlo_A1fw_HS2 | | BS02544A | RNA | 0/9 | 9 | 4.9 | 24,289 | NA | NA |
| 2 | C1fw_GGlo_A1rv_HS2 | | BS02566A | DNA | 9/1 | 9 | 18.8 | 62,474 | NA | NA |
| 2 | C1fw_GGlo_A1rv_HS2 | | BS02567A | DNA | 10/1 | 10 | 17.7 | 64,167 | NA | NA |
| 2 | C1fw_GGlo_A1rv_HS2 | | BS02611A | iPCR | 2/0 | NA | 12.7 | 21,508 | SE | 10,470 |
| 2 | C1fw_GGlo_A1rv_HS2 | | BS02611C | iPCR | 2/0 | NA | 21.7 | 24,583 | SE | 12,916 |
| 2 | C1fw_GGlo_A1rv_HS2 | | BS02612A | iPCR | 1/1 | NA | 11.0 | 32,938 | SE | 19,605 |
| 2 | C1fw_GGlo_A1rv_HS2 | | BS02545A | RNA | 2/10 | 12 | 4.5 | 36,846 | NA | NA |
| 2 | C1fw_GGlo_A1rv_HS2 | | BS02546A | RNA | 1/10 | 11 | 4.5 | 32,122 | NA | NA |

| Ex-peri-ment | Reporter Construct | Trans-fec-tion | Sample # | Type | Trim (R1/R2) | UMI length | Total read pairs (M) | Unique BC | Map-ping | Unique sites |
|---|---|---|---|---|---|---|---|---|---|---|
| 3 | GGlo | | BS02951A | DNA | 10/1 | 10 | 25.4 | 73,676 | NA | NA |
| 3 | GGlo | | BS03156A | DNA | 8/2 | 8 | 18.0 | 24,033 | NA | NA |
| 3 | GGlo | | BS03128A | iPCR | 2/1 | NA | 32.0 | 49,281 | SE | 31,838 |
| 3 | GGlo | | BS03128B | iPCR | 0/0 | NA | 1.1 | 47,228 | SE | 29,755 |
| 3 | GGlo | | BS03123A | RNA | 0/8 | 8 | 39.8 | 102,093 | NA | NA |
| 3 | GGlo | | BS03123B | RNA | 0/10 | 10 | 7.5 | 46,241 | NA | NA |
| 3 | GGlo | | BS03123C | RNA | 1/9 | 10 | 6.1 | 17,803 | NA | NA |
| 3 | GGlo | | BS03123D | RNA | 2/8 | 10 | 7.5 | 18,573 | NA | NA |
| 3 | GGlo_HS2 | | BS02953A | DNA | 8/1 | 8 | 26.7 | 33,430 | NA | NA |
| 3 | GGlo_HS2 | | BS03126A | iPCR | 0/1 | NA | 18.9 | 21,267 | SE | 14,515 |
| 3 | GGlo_HS2 | | BS03126B | iPCR | 1/0 | NA | 0.3 | 14,329 | SE | 9,794 |
| 3 | GGlo_HS2 | | BS03126C | iPCR | 0/1 | NA | 2.0 | 17,570 | SE | 12,713 |
| 3 | GGlo_HS2 | | BS03124A | RNA | 1/9 | 10 | 46.8 | 68,538 | NA | NA |
| 3 | GGlo_HS2 | | BS03124B | RNA | 0/8 | 8 | 8.0 | 55,196 | NA | NA |
| 3 | GGlo_HS2 | | BS03124C | RNA | 1/10 | 11 | 8.1 | 35,074 | NA | NA |
| 3 | GGlo_HS2 | | BS03124D | RNA | 2/9 | 11 | 9.5 | 36,509 | NA | NA |
| 3 | C1fw_GGlo_A1rv_HS2 | | BS02955A | DNA | 9/2 | 9 | 25.5 | 62,203 | NA | NA |
| 3 | C1fw_GGlo_A1rv_HS2 | | BS03158A | DNA | 10/0 | 10 | 19.4 | 36,871 | NA | NA |
| 3 | C1fw_GGlo_A1rv_HS2 | | BS03097A | iPCR | 0/1 | NA | 6.8 | 23,907 | SE | 13,125 |
| 3 | C1fw_GGlo_A1rv_HS2 | | BS03097B | iPCR | 1/0 | NA | 11.3 | 10,125 | SE | 5,307 |
| 3 | C1fw_GGlo_A1rv_HS2 | | BS03097I | iPCR | 0/1 | NA | 5.2 | 22,788 | SE | 13,989 |
| 3 | C1fw_GGlo_A1rv_HS2 | | BS03097J | iPCR | 0/1 | NA | 4.8 | 22,452 | SE | 13,156 |
| 3 | C1fw_GGlo_A1rv_HS2 | | BS03097K | iPCR | 0/1 | NA | 7.7 | 23,659 | SE | 12,650 |
| 3 | C1fw_GGlo_A1rv_HS2 | | BS03097L | iPCR | 0/1 | NA | 0.7 | 22,285 | SE | 13,337 |
| 3 | C1fw_GGlo_A1rv_HS2 | | BS03125A | RNA | 2/10 | 12 | 45.4 | 82,214 | NA | NA |
| 3 | C1fw_GGlo_A1rv_HS2 | | BS03125B | RNA | 0/9 | 9 | 5.6 | 45,788 | NA | NA |
| 3 | C1fw_GGlo_A1rv_HS2 | | BS03125C | RNA | 1/8 | 9 | 6.1 | 33,226 | NA | NA |
| 3 | C1fw_GGlo_A1rv_HS2 | | BS03125D | RNA | 2/10 | 12 | 7.0 | 30,931 | NA | NA |
| 4 | C1fw_GGlo_A1rv_HS2 | | BS03159A | DNA | 8/1 | 8 | 27.3 | 17,411 | NA | NA |
| 4 | C1fw_GGlo_A1rv_HS2 | | BS03160A | DNA | 9/0 | 9 | 33.2 | 44,490 | NA | NA |
| 4 | C1fw_GGlo_A1rv_HS2 | | BS03127A | iPCR | 1/0 | NA | 37.2 | 51,845 | SE | 35,813 |
| 4 | C1fw_GGlo_A1rv_HS2 | | BS03127B | iPCR | 0/0 | NA | 7.0 | 55,267 | SE | 37,733 |
| 4 | C1fw_GGlo_A1rv_HS2 | | BS03127C | iPCR | 2/1 | NA | 3.4 | 55,296 | SE | 37,909 |
| 4 | C1fw_GGlo_A1rv_HS2 | | BS03129A | iPCR | 0/0 | NA | 35.7 | 62,736 | SE | 42,308 |
| 4 | C1fw_GGlo_A1rv_HS2 | | BS03129B | iPCR | 1/1 | NA | 18.9 | 65,142 | SE | 42,565 |
| 4 | C1fw_GGlo_A1rv_HS2 | | BS03129C | iPCR | 1/1 | NA | 3.6 | 58,705 | SE | 39,682 |
| 4 | C1fw_GGlo_A1rv_HS2 | | BS03079A | RNA | 0/8 | 8 | 31.2 | 77,247 | NA | NA |
| 4 | C1fw_GGlo_A1rv_HS2 | | BS03196A | RNA | 0/8 | 8 | 16.2 | 66,379 | NA | NA |
| 4 | C1fw_GGlo_A1rv_HS2 | | BS03040B | 10xRNA | 0/0 | 12 | 39.3 | 18,661 | NA | NA |
| 4 | C1fw_GGlo_A1rv_HS2 | | BS03091A | 10xRNA | 0/0 | 12 | 32.7 | 18,153 | NA | NA |
| 4 | C1fw_GGlo_A1rv_HS2 | | BS03092A | 10xRNA | 0/0 | 12 | 26.3 | 16,642 | NA | NA |
| 4 | C1fw_GGlo_A1rv_HS2 | | BS03093A | 10xRNA | 0/0 | 12 | 15.4 | 14,621 | NA | NA |
| 4 | C1fw_GGlo_A1rv_HS2 | | BS03094A | 10xRNA | 0/0 | 12 | 29.8 | 18,301 | NA | NA |
| 4 | C1fw_GGlo_A1rv_HS2 | | BS03095A | 10xRNA | 0/0 | 12 | 36.0 | 19,198 | NA | NA |
| 5 | GGlo_HS2 | A | BS07222A | iPCR | 0/0 | NA | 5.1 | 47,683 | PE | 30,270 |
| 5 | GGlo_HS2 | A | BS07223A | iPCR | 1/1 | NA | 6.4 | 48,145 | PE | 29,938 |
| 5 | GGlo_HS2 | A | BS07381A | iPCR | 0/0 | NA | 5.0 | 67,184 | PE | 47,720 |
| 5 | GGlo_HS2 | A | BS07583A | iPCR | 0/0 | NA | 4.2 | 38,430 | PE | 26,364 |
| 5 | GGlo_HS2 | A | BS07587A | iPCR | 1/0 | NA | 6.4 | 38,146 | PE | 24,596 |
| 5 | C1CoreFw_GGlo_A1CoreFw_HS2 | B | BS07224A | iPCR | 2/0 | NA | 8.4 | 45,655 | PE | 27,172 |
| 5 | C1CoreFw_GGlo_A1CoreFw_HS2 | B | BS07225A | iPCR | 0/1 | NA | 4.9 | 47,475 | PE | 29,615 |
| 5 | C1CoreFw_GGlo_A1CoreFw_HS2 | B | BS07382A | iPCR | 1/1 | NA | 7.4 | 63,607 | PE | 43,027 |
| 5 | C1CoreFw_GGlo_A1CoreFw_HS2 | B | BS07584A | iPCR | 1/1 | NA | 5.3 | 37,811 | PE | 24,370 |
| 5 | C1CoreFw_GGlo_A1CoreFw_HS2 | B | BS07588A | iPCR | 2/1 | NA | 6.6 | 40,357 | PE | 25,466 |
| 5 | C1CoreFw_GGlo_A1CoreFw_HS2 | C | BS07226A | iPCR | 1/0 | NA | 6.6 | 45,233 | PE | 26,291 |
| 5 | C1CoreFw_GGlo_A1CoreFw_HS2 | C | BS07227A | iPCR | 2/1 | NA | 7.2 | 43,690 | PE | 23,970 |
| 5 | C1CoreFw_GGlo_A1CoreFw_HS2 | C | BS07383A | iPCR | 2/0 | NA | 9.2 | 66,322 | PE | 43,441 |
| 5 | C1CoreFw_GGlo_A1CoreFw_HS2 | C | BS07585A | iPCR | 2/0 | NA | 5.9 | 41,529 | PE | 27,038 |
| 5 | C1CoreFw_GGlo_A1CoreFw_HS2 | C | BS07589A | iPCR | 0/0 | NA | 4.5 | 40,809 | PE | 26,958 |
| 5 | C1CoreFw_GGlo_HS2_A1CoreFw | D | BS07228A | iPCR | 0/0 | NA | 4.9 | 30,767 | PE | 18,976 |
| 5 | C1CoreFw_GGlo_HS2_A1CoreFw | D | BS07229A | iPCR | 1/1 | NA | 6.8 | 31,771 | PE | 19,148 |
| 5 | C1CoreFw_GGlo_HS2_A1CoreFw | D | BS07384A | iPCR | 0/1 | NA | 5.1 | 42,841 | PE | 30,026 |
| 5 | C1CoreFw_GGlo_HS2_A1CoreFw | D | BS07586A | iPCR | 0/1 | NA | 4.2 | 29,283 | PE | 20,257 |
| 5 | C1CoreFw_GGlo_HS2_A1CoreFw | D | BS07590A | iPCR | 1/1 | NA | 6.3 | 29,076 | PE | 19,631 |
| 5 | GGlo_HS2 | A | BS07206A | DNA | 8/0 | 8 | 19.3 | 52,515 | NA | NA |
| 5 | GGlo_HS2 | A | BS07207A | DNA | 9/1 | 9 | 17.2 | 38,760 | NA | NA |
| 5 | GGlo_HS2 | A | BS07373A | DNA | 8/0 | 8 | 19.6 | 76,513 | NA | NA |
| 5 | GGlo_HS2 | A | BS07567A | DNA | 8/0 | 8 | 16.0 | 46,023 | NA | NA |

| Ex-peri-ment | Reporter Construct | Trans-fec-tion | Sample # | Type | Trim (R1/R2) | UMI length | Total read pairs (M) | Unique BC | Map-ping | Unique sites |
|---|---|---|---|---|---|---|---|---|---|---|
| 5 | GGlo_HS2 | A | BS07571A | DNA | 9/0 | 9 | 23.3 | 48,033 | NA | NA |
| 5 | C1CoreFw_GGlo_A1CoreFw_HS2 | B | BS07208A | DNA | 10/2 | 10 | 15.7 | 31,704 | NA | NA |
| 5 | C1CoreFw_GGlo_A1CoreFw_HS2 | B | BS07209A | DNA | 8/2 | 8 | 17.3 | 30,861 | NA | NA |
| 5 | C1CoreFw_GGlo_A1CoreFw_HS2 | B | BS07374A | DNA | 9/2 | 9 | 17.1 | 64,026 | NA | NA |
| 5 | C1CoreFw_GGlo_A1CoreFw_HS2 | B | BS07568A | DNA | 9/1 | 9 | 15.4 | 38,288 | NA | NA |
| 5 | C1CoreFw_GGlo_A1CoreFw_HS2 | B | BS07572A | DNA | 10/1 | 10 | 20.7 | 40,668 | NA | NA |
| 5 | C1CoreFw_GGlo_A1CoreFw_HS2 | C | BS07210A | DNA | 9/0 | 9 | 16.4 | 50,833 | NA | NA |
| 5 | C1CoreFw_GGlo_A1CoreFw_HS2 | C | BS07211A | DNA | 10/1 | 10 | 17.4 | 25,135 | NA | NA |
| 5 | C1CoreFw_GGlo_A1CoreFw_HS2 | C | BS07375A | DNA | 10/1 | 10 | 17.1 | 59,315 | NA | NA |
| 5 | C1CoreFw_GGlo_A1CoreFw_HS2 | C | BS07569A | DNA | 10/2 | 10 | 15.2 | 40,267 | NA | NA |
| 5 | C1CoreFw_GGlo_A1CoreFw_HS2 | C | BS07573A | DNA | 8/1 | 8 | 21.9 | 37,500 | NA | NA |
| 5 | C1CoreFw_GGlo_HS2_A1CoreFw | D | BS07212A | DNA | 8/1 | 8 | 18.1 | 17,813 | NA | NA |
| 5 | C1CoreFw_GGlo_HS2_A1CoreFw | D | BS07213A | DNA | 9/2 | 9 | 17.1 | 29,070 | NA | NA |
| 5 | C1CoreFw_GGlo_HS2_A1CoreFw | D | BS07376A | DNA | 9/2 | 9 | 13.8 | 42,187 | NA | NA |
| 5 | C1CoreFw_GGlo_HS2_A1CoreFw | D | BS07570A | DNA | 8/2 | 8 | 15.5 | 27,218 | NA | NA |
| 5 | C1CoreFw_GGlo_HS2_A1CoreFw | D | BS07574A | DNA | 9/2 | 9 | 17.7 | 26,428 | NA | NA |
| 5 | GGlo_HS2 | A | BS07214A | RNA | 0/8 | 8 | 14.2 | 83,166 | NA | NA |
| 5 | GGlo_HS2 | A | BS07215A | RNA | 1/9 | 10 | 13.6 | 59,645 | NA | NA |
| 5 | GGlo_HS2 | A | BS07377A | RNA | 0/10 | 10 | 13.5 | 94,915 | NA | NA |
| 5 | GGlo_HS2 | A | BS07575A | RNA | 0/8 | 8 | 13.9 | 68,189 | NA | NA |
| 5 | GGlo_HS2 | A | BS07579A | RNA | 1/10 | 11 | 12.9 | 56,750 | NA | NA |
| 5 | C1CoreFw_GGlo_A1CoreFw_HS2 | B | BS07216A | RNA | 2/10 | 12 | 14.5 | 49,094 | NA | NA |
| 5 | C1CoreFw_GGlo_A1CoreFw_HS2 | B | BS07217A | RNA | 0/9 | 9 | 14.8 | 67,064 | NA | NA |
| 5 | C1CoreFw_GGlo_A1CoreFw_HS2 | B | BS07378A | RNA | 1/8 | 9 | 15.0 | 72,061 | NA | NA |
| 5 | C1CoreFw_GGlo_A1CoreFw_HS2 | B | BS07576A | RNA | 1/9 | 10 | 14.1 | 50,898 | NA | NA |
| 5 | C1CoreFw_GGlo_A1CoreFw_HS2 | B | BS07581A | RNA | 0/10 | 10 | 13.2 | 64,310 | NA | NA |
| 5 | C1CoreFw_GGlo_A1CoreFw_HS2 | C | BS07218A | RNA | 1/10 | 11 | 14.2 | 47,994 | NA | NA |
| 5 | C1CoreFw_GGlo_A1CoreFw_HS2 | C | BS07219A | RNA | 2/8 | 10 | 13.8 | 57,832 | NA | NA |
| 5 | C1CoreFw_GGlo_A1CoreFw_HS2 | C | BS07379A | RNA | 2/9 | 11 | 15.7 | 90,459 | NA | NA |
| 5 | C1CoreFw_GGlo_A1CoreFw_HS2 | C | BS07577A | RNA | 2/10 | 12 | 15.6 | 64,030 | NA | NA |
| 5 | C1CoreFw_GGlo_A1CoreFw_HS2 | C | BS07580A | RNA | 2/8 | 10 | 17.3 | 66,033 | NA | NA |
| 5 | C1CoreFw_GGlo_HS2_A1CoreFw | D | BS07220A | RNA | 0/10 | 10 | 12.6 | 47,992 | NA | NA |
| 5 | C1CoreFw_GGlo_HS2_A1CoreFw | D | BS07221A | RNA | 1/8 | 9 | 14.5 | 32,527 | NA | NA |
| 5 | C1CoreFw_GGlo_HS2_A1CoreFw | D | BS07380A | RNA | 1/10 | 11 | 16.0 | 55,183 | NA | NA |
| 5 | C1CoreFw_GGlo_HS2_A1CoreFw | D | BS07578A | RNA | 0/9 | 9 | 12.8 | 41,321 | NA | NA |
| 5 | C1CoreFw_GGlo_HS2_A1CoreFw | D | BS07582A | RNA | 1/8 | 9 | 15.9 | 40,467 | NA | NA |
| 5 | Pool1 | Pool1 | BS07509A | 10xRNA | 0/0 | 12 | 17.0 | 88,525 | NA | NA |
| 5 | Pool1 | Pool1 | BS07623A | 10xRNA | 0/0 | 12 | 17.3 | 85,901 | NA | NA |
| 5 | Pool1 | Pool1 | BS07700A | 10xRNA | 0/0 | 12 | 6.4 | 62,377 | NA | NA |
| 5 | Pool1 | Pool1 | BS07704A | 10xRNA | 0/0 | 12 | 5.3 | 69,276 | NA | NA |
| 5 | Pool2 | Pool2 | BS07510A | 10xRNA | 0/0 | 12 | 18.6 | 83,620 | NA | NA |
| 5 | Pool2 | Pool2 | BS07624A | 10xRNA | 0/0 | 12 | 18.2 | 80,386 | NA | NA |
| 5 | Pool2 | Pool2 | BS07701A | 10xRNA | 0/0 | 12 | 6.2 | 58,564 | NA | NA |
| 5 | Pool2 | Pool2 | BS07705A | 10xRNA | 0/0 | 12 | 6.2 | 65,772 | NA | NA |
| 5 | Pool3 | Pool3 | BS07511A | 10xRNA | 0/0 | 12 | 21.0 | 87,274 | NA | NA |
| 5 | Pool3 | Pool3 | BS07625A | 10xRNA | 0/0 | 12 | 24.0 | 87,971 | NA | NA |
| 5 | Pool3 | Pool3 | BS07702A | 10xRNA | 0/0 | 12 | 6.8 | 64,088 | NA | NA |
| 5 | Pool3 | Pool3 | BS07706A | 10xRNA | 0/0 | 12 | 5.9 | 67,286 | NA | NA |
| 5 | Pool4 | Pool4 | BS07512A | 10xRNA | 0/0 | 12 | 18.7 | 83,824 | NA | NA |
| 5 | Pool4 | Pool4 | BS07626A | 10xRNA | 0/0 | 12 | 22.2 | 84,701 | NA | NA |
| 5 | Pool4 | Pool4 | BS07703A | 10xRNA | 0/0 | 12 | 6.8 | 52,790 | NA | NA |
| 5 | Pool4 | Pool4 | BS07707A | 10xRNA | 0/0 | 12 | 4.7 | 59,714 | NA | NA |

**Supplemental Table S3. Summaries of analyzed barcodes by transfection.**
Shown are counts of BCs analyzed for the individual transfections in each experiment.

| Experiment | Sample | DNA | RNA | iPCR | Analyzed sites |
|---|---|---|---|---|---|
| Experiment 1 | GGlo | 178,306 | 90,165 | 95,902 | 66,742 |
| Experiment 1 | GGlo+HS2 | 129,407 | 90,267 | 36,541 | 28,305 |
| Experiment 1 | Ins+GGlo+Ins+HS2 | 59,959 | 35,108 | 21,974 | 18,148 |
| Experiment 1 | Ins+GGlo+Ins+HS2 | 85,449 | 76,983 | 39,520 | 28,446 |
| Experiment 2 | GGlo | 86,288 | 58,726 | 29,481 | 25,930 |
| Experiment 2 | GGlo+HS2 | 97,499 | 78,318 | 25,647 | 21,572 |
| Experiment 2 | Ins+GGlo+Ins+HS2 | 85,363 | 40,729 | 14,939 | 12,390 |
| Experiment 2 | Ins+GGlo+Ins+HS2 | 90,316 | 52,751 | 29,988 | 22,968 |
| Experiment 3 | GGlo | 85,691 | 113,862 | 44,670 | 28,107 |
| Experiment 3 | GGlo+HS2 | 33,430 | 85,129 | 18,392 | 11,974 |
| Experiment 3 | Ins+GGlo+Ins+HS2 | 69,569 | 95,375 | 22,995 | 18,469 |
| Experiment 4 | Ins+GGlo+Ins+HS2 | 49,923 | 89,502 | 55,045 | 30,935 |

**Supplemental Table S4. Summary of 3' 10x libraries.**
Shown are 10x scRNA-seq libraries, sequencing statistics, and mapping statistics.

| Experiment | Pool | # cells loaded | Sample # | # Reads | Estimated # Cells | Median Reads per cell | Median Genes per cell | Median UMIs per Cell |
|---|---|---|---|---|---|---|---|---|
| Experiment 5 | Pool1 | 27,094 | BS07468 | 2,220,021,482 | 25,480 | 87,128 | 3,610 | 10,760 |
| Experiment 5 | Pool2 | 25,048 | BS07469 | 1,389,186,887 | 11,832 | 117,409 | 5,951 | 35,060 |
| Experiment 5 | Pool3 | 28,226 | BS07470 | 1,499,082,944 | 13,086 | 114,556 | 5,975 | 34,896 |
| Experiment 5 | Pool4 | 19,632 | BS07471 | 1,379,307,019 | 12,024 | 114,713 | 5,722 | 32,213 |

### Supplemental Table S5. PCR primer and DNA fragment sequences.

The applicable Library Type is indicated where relevant.

| Primer/DNA Fragment Name | Sequence | Library Type |
|---|---|---|
| P5_barcode_0N | TTTCCCTACACGACGCTCTTCCGATCTCCTTTCTAGGCAATTAGG | RNA, iPCR |
| P5_barcode_1N | TTTCCCTACACGACGCTCTTCCGATCTNCCTTTCTAGGCAATTAGG | RNA, iPCR |
| P5_barcode_2N | TTTCCCTACACGACGCTCTTCCGATCTNNCCTTTCTAGGCAATTAGG | RNA, iPCR |
| P7_barcode_0N | CTGGAGTTCAGACGTGTGCTCTTCCGATCTCCTTTCTAGGCAATTAGG | DNA |
| P7_barcode_1N | CTGGAGTTCAGACGTGTGCTCTTCCGATCTNCCTTTCTAGGCAATTAGG | DNA |
| P7_barcode_2N | CTGGAGTTCAGACGTGTGCTCTTCCGATCTNNCCTTTCTAGGCAATTAGG | DNA |
| P7_ITR_0N | CTGGAGTTCAGACGTGTGCTCTTCCGATCTCTTCCGACTTCAACTGTA | iPCR |
| P7_ITR_1N | CTGGAGTTCAGACGTGTGCTCTTCCGATCTNCTTCCGACTTCAACTGTA | iPCR |
| P7_ITR_2N | CTGGAGTTCAGACGTGTGCTCTTCCGATCTNNCTTCCGACTTCAACTGTA | iPCR |
| P5_plasmid_8N | TTTCCCTACACGACGCTCTTCCGATCTNNNNNNNNAGCTGCACAGCAACAC | DNA |
| P5_plasmid_9N | TTTCCCTACACGACGCTCTTCCGATCTNNNNNNNNNAGCTGCACAGCAACAC | DNA |
| P5_plasmid_10N | TTTCCCTACACGACGCTCTTCCGATCTNNNNNNNNNNAGCTGCACAGCAACAC | DNA |
| P7_plasmid_8N | CTGGAGTTCAGACGTGTGCTCTTCCGATCTNNNNNNNNAGCTGCACAGCAACAC | RNA |
| P7_plasmid_9N | CTGGAGTTCAGACGTGTGCTCTTCCGATCTNNNNNNNNNAGCTGCACAGCAACAC | RNA |
| P7_plasmid_10N | CTGGAGTTCAGACGTGTGCTCTTCCGATCTNNNNNNNNNNAGCTGCACAGCAACAC | RNA |
| P7_10xSB-barcodeV2_0N | CTGGAGTTCAGACGTGTGCTCTTCCGATCTTTGGACAAAGATCCCACAACTAGC | 10xRNA |
| P7_halfsite_S | CTGGAGTTCAGACGTGTGCT | RNA |
| P5_halfsite_Short | TTTCCCTACACGACGCTCTT | DNA, RNA |
| P5_halfsite | TTTCCCTACACGACGCTCTTCCGATCT | 10xRNA |
| P7_amplicon_N704 | CAAGCAGAAGACGGCATACGAGATGCTCAGGAGTGACTGGAGTTCAGACGTGTGCT | DNA, RNA, iPCR |
| P7_amplicon_N705 | CAAGCAGAAGACGGCATACGAGATAGGAGTCCGTGACTGGAGTTCAGACGTGTGCT | DNA, RNA, iPCR |
| P7_amplicon_N706 | CAAGCAGAAGACGGCATACGAGATCATGCCTAGTGACTGGAGTTCAGACGTGTGCT | DNA, RNA, iPCR |
| P7_amplicon_N707 | CAAGCAGAAGACGGCATACGAGATGTAGAGAGGTGACTGGAGTTCAGACGTGTGCT | DNA, RNA, iPCR |
| P7_amplicon_N710 | CAAGCAGAAGACGGCATACGAGATCAGCCTCGGTGACTGGAGTTCAGACGTGTGCT | DNA, RNA, iPCR |
| P7_amplicon_N711 | CAAGCAGAAGACGGCATACGAGATTGCCTCTTGTGACTGGAGTTCAGACGTGTGCT | DNA, RNA, iPCR |
| P7_amplicon_N712 | CAAGCAGAAGACGGCATACGAGATTCCTCTACGTGACTGGAGTTCAGACGTGTGCT | DNA, RNA, iPCR |
| P7_amplicon_N714 | CAAGCAGAAGACGGCATACGAGATTCATGAGCGTGACTGGAGTTCAGACGTGTGCT | DNA, RNA, iPCR |
| P7_amplicon_N715 | CAAGCAGAAGACGGCATACGAGATCCTGAGATGTGACTGGAGTTCAGACGTGTGCT | DNA, RNA, iPCR |
| P7_amplicon_N716 | CAAGCAGAAGACGGCATACGAGATTAGCGAGTGTGACTGGAGTTCAGACGTGTGCT | DNA, RNA, iPCR |
| P7_amplicon_N718 | CAAGCAGAAGACGGCATACGAGATGTAGCTCCGTGACTGGAGTTCAGACGTGTGCT | DNA, RNA, iPCR |
| P7_amplicon_N719 | CAAGCAGAAGACGGCATACGAGATTACTACGCGTGACTGGAGTTCAGACGTGTGCT | DNA, RNA, iPCR |
| P7_amplicon_N720 | CAAGCAGAAGACGGCATACGAGATAGGCTCCGGTGACTGGAGTTCAGACGTGTGCT | DNA, RNA, iPCR |
| P5_amplicon_S502 | AATGATACGGCGACCACCGAGATCTACACCTCTCTATACACTCTTTCCCTACACGACGCTCTT | DNA, RNA, iPCR |
| P5_amplicon_S503 | AATGATACGGCGACCACCGAGATCTACACTATCCTCTACACTCTTTCCCTACACGACGCTCTT | DNA, RNA, iPCR |
| P5_amplicon_S505 | AATGATACGGCGACCACCGAGATCTACACGTAAGGAGACACTCTTTCCCTACACGACGCTCTT | DNA, RNA, iPCR |
| P5_amplicon_S506 | AATGATACGGCGACCACCGAGATCTACACACTGCATAACACTCTTTCCCTACACGACGCTCTT | DNA, RNA, iPCR |
| P5_amplicon_S507 | AATGATACGGCGACCACCGAGATCTACACAAGGAGTAACACTCTTTCCCTACACGACGCTCTT | DNA, RNA, iPCR |
| P5_amplicon_S508 | AATGATACGGCGACCACCGAGATCTACACCTAAGCCTACACTCTTTCCCTACACGACGCTCTT | DNA, RNA, iPCR |
| P5_amplicon_S510 | AATGATACGGCGACCACCGAGATCTACACCGTCTAATACACTCTTTCCCTACACGACGCTCTT | DNA, RNA, iPCR |
| P5_amplicon_S511 | AATGATACGGCGACCACCGAGATCTACACTCTCTCCGACACTCTTTCCCTACACGACGCTCTT | DNA, RNA, iPCR |
| P5_amplicon_S513 | AATGATACGGCGACCACCGAGATCTACACTCGACTAGACACTCTTTCCCTACACGACGCTCTT | DNA, RNA, iPCR |
| P5_amplicon_S515 | AATGATACGGCGACCACCGAGATCTACACTTCTAGCTACACTCTTTCCCTACACGACGCTCTT | DNA, RNA, iPCR |
| P5_amplicon_S516 | AATGATACGGCGACCACCGAGATCTACACCCTAGAGTACACTCTTTCCCTACACGACGCTCTT | DNA, RNA, iPCR |
| P5_amplicon_S517 | AATGATACGGCGACCACCGAGATCTACACGCGTAAGAACACTCTTTCCCTACACGACGCTCTT | DNA, RNA, iPCR |
| P5_amplicon_S518 | AATGATACGGCGACCACCGAGATCTACACCTATTAAGACACTCTTTCCCTACACGACGCTCTT | DNA, RNA, iPCR |
| P5_amplicon_S520 | AATGATACGGCGACCACCGAGATCTACACAAGGCTATACACTCTTTCCCTACACGACGCTCTT | DNA, RNA, iPCR |
| P5_amplicon_S521 | AATGATACGGCGACCACCGAGATCTACACGAGCCTTAACACTCTTTCCCTACACGACGCTCTT | DNA, RNA, iPCR |
| P5_amplicon_S522 | AATGATACGGCGACCACCGAGATCTACACTTATGCGAACACTCTTTCCCTACACGACGCTCTT | DNA, RNA, iPCR |
| Gibson BC4 | CcCGAGCTCGATGAGTTTGGACAAA-gatcCCACAACTAGcnnnnnnnnnnnnnnnCCTAATTGCCTAGAAAGGAGCAGACGATATGGC | n/a |
| GGlo | TAAAGGGAAGAATAAATTAGAGAAAAACTGGAATGACTGAATCGGAACAAGGCAAAGGC-TATAAAAAAAATTAAGCAGCAGTATCCTCTTGGGGGCCCCTTCCCCACACTATCTCAATGCAAATATCTGTCTGAAACGGTCCCTGGCTAAACTCCACCCATGGGTTGGCCAGCCTTGCCTTGACCAA-TAGCCTTGACAAGGCAAACTTGACCAATAGTCTTAGAGTATCCAGTGAGGCCAGGGGCCGGCGGCTGGCTAGGGATGAAGAATAAAAGGAAGCACCCTTCAGCAGTTCCACACACTCGCTTCTGGAAC-GTCTGAGGTTATCAATAAGGACCTGCAGCCAAGCTTGAGCTTCACGCTGCCGCAAGCACTCAGGGCGCAAGGGCTGCTAAAGGAAGCGGAACACGTAGAAAGCCAGTCCGCAGAAAC-GGTGCTGACCCCGGATGAATGTCAGCTACTGGGCTATCTGGACAAGGGAAAACGCAAGCGCAAAGAGAAAGCAGGTAGCTTGCAGTGGGCTTACATGGCGATAGCTA-GACTGGGCGGTTTTATGGACAGCAAGCGAACCGGAATTGCCAGCTGGGGCGCCCTCTGGTAAGGTTGGGAAGCCCTGCAAAGTAAACTGGATGGCTTTCTTGCCGCCAAGGATCTGATGGCGCAGGG-GATCAAGATCTGATCAAGAGACAGGATGAGGATCGTTTCGCA | n/a |

| Primer/DNA Fragment Name | Sequence | Library Type |
|---|---|---|
| HS2 | ACCGGTTGTTTCCTTATCTGACCTGCTTTAACTGGGTAAGCTTATGAAAAGTCTTGTG-<br>TAGAAGAGAAAGGGATAACAGCCTGTGCTAAATGAGGAAGCTGCCTTGCCTGTTCCTGCTCAGTGGG<br>GTTTCTGGCTATACTACATCAACTGAGTCAGTGCATCTTGTCTAGTTCCCACAC-<br>CTTTCTCTGAACAGAGAGAGTAAAGGGCTCAATAAGAAAAATACAGTTTATGGTCTGTACGTTGTATTA<br>CACGTCATCCCTTACTTTCTAAAAGGCATCTTCACTGAGAAAGACATGGATTTCTAAC-<br>CTTACAATCTCTACAAAGTTCAGAATACATAATAATCTTGAATTATGATTAACTGTTAGTTTTGACCAGG<br>TCTTCTGGCAGACAGGTCACATGTGTTAGTATCACTTATTCCTCAAGTGTTGATGTTAG-<br>TGTCAGCATATTACCGATGTTCCACAAACATTTCCTGAATGACTGTTAAACTTCCTACACATTAACGAG<br>CCTCTGCATTTTTTTCCAGCTTCCATCTATGATTTAAGTAAACTCTAGTTTTCCAC-<br>TTCTTCATATTCTCTCTCTACATCTCAATTATTGCAGTACCACTGTCCAAGGGCAGAGGAGGTTAGCT<br>GGGCCCAGGCGGAGTCAATTCTCTACTCCCCACCCTGTGGGTGTGTTCAGCCTT-<br>GTGAGCCAGCATCAGGCTTGAGCACAGCAGTGCTGAGTCATGCTGAGTCATGCTGAGGCTTAGGGT<br>GTGTGGCCAGATGTTTTCAGCTGTGAGTCATCAGTGCTATCTGGGTCTCTAGGAG-<br>GAAGTCCACAGGGAAGGTGAAAAGAAAATAAGTTTGCTCCCTGAAGAAAACATTACTTACACCAGCAT<br>TACAATGAAAAGGGGACCCTGCCTTGCTGTGTGACATAACCTAGAATATTTTATTTTCTAG-<br>TTAAAAATTACTTCTCATCATCTTTATTAGAAGTTATAACAAATCATTTTTTAGAAATGTAAAGTTTATTT<br>AGACTCATACCTAGAAGCAACAGAATCACACATTTTAGTGTACACACACACACACACAC-<br>GCACACACACACACATGACTTTTGTTCATAAACTATGCAATTGTTAATGAAATGCTATTTGGAATGGGC<br>TTGAATTCGTATTCC | n/a |
| A1 insulator | CTGAACAGGTTGACTATTAATTGTGTCTGCTTGATGTGGACACCAGGTGGCGCTG-<br>GACATCAGATTTGGAGAGGCAGTTGTCTAGGGAACCGGGCTCTGTGCCAGCGCAGGAGGCAGGCTG<br>GCTCTCCTATTCCAGGGATGCTCATCCAGGAAGGAAAGGTTGCATGCTGGACACACTAACCT | n/a |
| C1 insulator | AGGAAAAGCCCCAGAGGATGTCCCCCCGCGTTCATACCCTAAGAGAAGATGCAACAG-<br>GAACAAGAAATTGCACAACAGAGCTCTGTGGACCATGAAAAGGCCCCTTCCTCCACAGTGTCCCTCC<br>GGCGCCCTCTACTGACAAAGCTTGATATCGTGCTTGCTGCAAAGAAGGGCTTAGAGTCCAG-<br>TCCATCAAGGAGCAGGCACTCACGGGGAAGTTGGAAGCTGAGAGGCAAGAAGAAACCAG | n/a |
| A1Core insulator | ATCTGATGTCCAGCGCCACCTGGTGTCCACATCAAGCAGACACAATTAATAGTC | n/a |
| C1Core insulator | AGTGTCCCTCCGGCGCCCTCTACTGACAAAGCTTGATATCGTGCTTGCTGCAAA | n/a |

**Supplemental Table S6. Plasmids.**
Summary of plasmids used and their derivation if applicable.

| Plasmid # | Plasmid Name | Starting Plasmid | Element(s) Removed | Synthetic DNA Inserted |
|---|---|---|---|---|
| pMH012 | pTREM-Bglo-GFP | pT2/LTR7-GFP (Addgene #62541) | LTR7 promoter | Human Beta-globin minimal promoter |
| pMH019 | pTR-GGlo-GFP | pMH012 | Human ß-globin minimal promoter | Gamma-globin promoter |
| pMH020 | pTR-GGlo-GFP-HS2 | pMH012 | n/a | Gamma-globin promoter, Mouse HS2 enhancer |
| pMH021 | pTR-GGlo-GFP-A1rv-HS2 | pMH012 | n/a | Gamma-globin promoter, Mouse HS2 enhancer, 3' A1rv insulator |
| pMH024 | pTR-A1fw-GGlo-GFP-A1fw-HS2 | pMH012 | n/a | Gamma-globin promoter, Mouse HS2 enhancer, 5' A1fw insulator, 3' A1fw insulator |
| pMH024B | pTR-A1fw-GGlo-PuroP2AGFP-A1fw-HS2 | pMH024 | n/a | Puromycin resistance, P2A sequence |
| pMH027 | pTR_GGlo_PuroP2A_GFP | pMH019 | n/a | Puromycin resistance, P2A sequence |
| pMH028 | pTR-GGlo-PuroP2AGFP-HS2 | pMH020 | n/a | Puromycin resistance, P2A sequence |
| pMH030 | pTR-GGlo-PuroP2AGFP-A1rv-HS2 | pMH021 | n/a | Puromycin resistance, P2A sequence |
| pMH032 | pTR-C1fw-GGlo-PuroP2AGFP-A1fw-HS2 | pMH024B | n/a | C1fw Insulator |
| pMH034 | pTR-C1fw-GGlo-PuroP2AGFP-A1rv-HS2 | pMH030 | n/a | C1fw Insulator |
| pMH035 | SB100X-mKate_v3 | pCMV(CAT)T7-SB100 (Addgene plasmid #34879) | n/a | mKate |
| pMH040 | pTR-A1CoreRV-GGlo-PuroP2AGFP-A1CoreFW-HS2 | pMH028 | n/a | A1CoreFW Insulator, A1CoreRV Insulator |
| pMH041 | pTR_C1CoreFW_GGlo_PuroP2AGFP_A1CoreFW_HS2 | pMH028 | n/a | A1CoreFW Insulator, C1CoreFW Insulator |
| pMH049 | pTR_C1fw_GGlo_PuroP2AGFP_A1fw_minusHS2 | pMH032 | Mouse HS2 Enhancer | n/a |
| pMH050 | pMH050_pTR_C1fw_PuroP2AGFP_A1fw_HS2_minusGglo | pMH032 | Gamma-globin promoter | n/a |
| pMH065 | pTR_C1CoreFW_GGlo_PuroP2AGFP_HS2_A1Corefw | pMH028 | n/a | A1CoreFW Insulator, C1CoreFW Insulator |

**Supplemental Data**

**Supplemental Data S1. Clonal inference analysis results for Experiment 4.**
Each line corresponds to a distinct clone. BCs and cellBCs contain a comma-separated list of the Reporter BCs and cellBCs assigned to that clone. clone contains a unique clone ID. count, total UMIs. nedges, number of reporter BC to cell BC. nBCs, number of unique reporter BCs. ncells, number of unique cellBCs.

**Supplemental Data S2. Clonal inference analysis results for Experiment 5.**
Format is the same as **Supplemental Data S2**, with an additional column transfection that indicates the inferred source transfection of the clone.

**Supplemental Data S3. Reporter integration analysis of gene perturbation.**
Each row represents a differential expression test between a reporter and gene. Reporter_chrom, Reporter_chromStart. Reporter_Class, and Reporter_BC record reporter genomic coordinates (hg38), reporter class, and reporter BC. TSS_chromStart, Gene_Symbol, and Ensembl_ID indicate the gene TSS position (hg38), gene symbol and ENSEMBL gene ID. DistToTSS indicates the distance from reporter insertion to gene TSS. Within_gene_body is a flag indicating whether the reporter lies within gene body. n, number of cells in the reporter clone. x0 and x1, average gene UMI count across unperturbed and perturbed cells (respectively). mu, overall gene average UMI count. xb, estimated gene average UMI count among perturbed cells. log2_fold_change, expression change in perturbed cells. z, perturbation z-score. p.value, significance of perturbation. q.value significance corrected for multiple testing.
